# Age-related changes in the neural processing of semantics, within and beyond the core semantic network

**DOI:** 10.1101/2023.05.05.539561

**Authors:** Wei Wu, Paul Hoffman

**Author notes:** Correspondence to: Dr. Paul Hoffman School of Philosophy, Psychology & Language Sciences, University of Edinburgh, 7 George Square, Edinburgh, EH8 9JZ, UK : Dr. Wei Wu School of Philosophy, Psychology & Language Sciences, University of Edinburgh, 7 George Square, Edinburgh, EH8 9JZ, UK or. Declarations of interest: none.

## Abstract

Ageing is associated with increases in functional activation, which have been interpreted either as compensatory responses to the higher task demands older people experience, or as neural dedifferentiation. Ageing is also characterised by a shift to greater reliance on prior knowledge and less on executive function, whose underlying neural mechanism is poorly understood. This pre-registered fMRI study investigated these questions within the domain of semantic cognition. To disentangle the compensation and dedifferentiation theories, we extracted activation signal in core verbal semantic regions, for young and older participants during semantic tasks. Verbal semantic processing relies heavily on left inferior frontal gyrus (IFG) but older people frequently show additional right IFG activation. We found that right IFG exhibited a similar linear activation-demand relationship as left IFG across age groups and semantic tasks, indicating that age-related over-recruitment of this region may be compensatory in nature. To answer the second question, we examined network-level activity and connectivity changes in semantic and non-semantic tasks. Older people showed more engagement of the default mode network (DMN) and less of the executive multiple demand network (MDN) aligning with their greater reserves of prior knowledge and declined executive control. In contrast, activation was age-invariant in regions contributing specifically to executive control of semantic processing. Older adults also showed a degraded ability to modulate MDN activation as a function of demand in the non-semantic task, but not in the semantic tasks. These findings provide a new perspective on the neural basis of semantic cognition in later life, and suggest that preservation of activation in specialised semantic networks may support preserved performance in this critical domain.

## Introduction

Healthy ageing is accompanied by reorganization in brain function; a key goal of cognitive neuroscience is to understand these reorganizations and their cognitive bases (Grady, 2012). While various forms of functional change have been investigated, a commonly reported pattern is that older adults exhibit a more bilateral pattern of activation in task-relevant regions (especially in prefrontal cortex) across a range of cognitive tasks (Berlingeri, Danelli, Bottini, Sberna, & Paulesu, 2013; Cabeza et al., 1997; Hoffman & Morcom, 2018; Spreng, Wojtowicz, & Grady, 2010). This effect has been termed Hemispheric Asymmetry Reduction in OLDer adults (HAROLD; Cabeza, 2002). Although many studies have reported this pattern, there is limited understanding of the mechanisms behind it. The current study investigated this question in the domain of semantic cognition. Networks related to language and semantic processing are usually characterized as left-lateralised, but smaller amounts of activation in right-hemisphere homologous regions are also frequently observed (Jackson, 2021; Rice, Lambon Ralph, & Hoffman, 2015). A recent meta-analysis found that this right-hemisphere activation was greater in older people, especially in prefrontal cortex (Hoffman & Morcom, 2018). The functional significance of the activation in the right homologue, and its increase in later life, is unclear. In the present study, we aimed to reveal the right hemisphere regions’ contribution to semantic processing across the life span, as well as understanding more general changes in neural correlates of semantic processing in the ageing brain.

The ongoing debate over age-related activation changes can be summarized into two major accounts - *compensation* versus *dedifferentiation*. Compensation refers to the recruitment of additional neural resources in response to heightened cognitive demands (Cabeza et al., 2018; D. C. Park & Reuter-Lorenz, 2009; Reuter-Lorenz & Cappell, 2008). On the compensation view, older people show a more bilateral pattern of activation when they find tasks more cognitively demanding than young people. In these situations, core task-relevant regions approach or exceed their processing limits and the contralateral hemisphere is recruited to provide additional support. In contrast, the dedifferentiation view proposes that neural activity becomes less selective and specific in older age, leading to superfluous neural activations that do not contribute to task processing (Li & Rieckmann, 2014; Logan, Sanders, Snyder, Morris, & Buckner, 2002; Morcom & Henson, 2018; Nyberg et al., 2014; D. C. Park et al., 2004). On this view, increased contralateral recruitment in old age does not support task performance and may even be maladaptive.

Several theories have been proposed in support of the compensation account. The Scaffolding Theory of Aging and Cognition (STAC) has described compensatory activation in older age as a scaffolding response, which protects cognitive performance by engaging alternative neural circuits that assist in processing (D. C. Park & Reuter-Lorenz, 2009). Moreover, STAC suggests that scaffolding is a dynamic process across the lifespan and is not limited to old age. For example, it can also help to characterize neural dynamics in young adults when they acquire novel skills (D. C. Park & Reuter-Lorenz, 2009; Reuter-Lorenz & Park, 2014). Another theory named CRUNCH (Compensation-Related Utilization of Neural Circuits Hypothesis) has drawn a detailed trajectory illustrating how neural activation changes with cognitive demands (Reuter-Lorenz & Cappell, 2008). According to CRUNCH, when a particular task is more demanding to older people than to young, the seniors need more neural resources to achieve the same cognitive goal. This leads to increased contralateral activation as a resource ceiling or activation plateau is approached in the dominant hemisphere. Crucially, the same demand-activation trajectory occurs in the junior brain but at higher levels of task difficulty, as young people usually perform tasks more efficiently than their older counterparts (Schneider-Garces et al., 2010). Thus, both theories propose that compensatory responses are present across age groups under conditions of increasing cognitive challenge. Greater contralateral activation is typically observed in older people because they usually find tasks more challenging than young people. If this position were reversed and young people were exposed to a task they found more challenging than older people, the same compensatory regions should show greater activation in the young group. We test this novel hypothesis in the present study.

The dedifferentiation account was largely built on cognitive ageing findings in domain-selective ventral occipitotemporal cortex (VOTC; Caramazza & Mahon, 2003; Downing, Chan, Peelen, Dodds, & Kanwisher, 2005). VOTC is a higher-level visual region that contains clusters showing selective responses to objects from certain domains (e.g., the fusiform face area to faces). Using a passive viewing paradigm, an early and influential study found that domain-selective effects in VOTC were reduced in older participants, and this was interpreted as less neural specialization in the ageing brain (D. C. Park et al., 2004). Similar age-related neural dedifferentiation in VOTC has been replicated in subsequent studies (J. Park et al., 2012; Voss et al., 2008), and some research has associated it with deteriorated performance in mnemonic object recognition tasks (Berron et al., 2018; but see Koen, Hauck, & Rugg, 2019). A computational model of dedifferentiation proposes that the senior brain has decreased neuromodulator availability (e.g., dopamine), which can jeopardize the fidelity of neural signalling and thus lead to less distinctiveness in neural representations (Li, Brehmer, Shing, Werkle-Bergner, & Lindenberger, 2006; Li & Rieckmann, 2014). It has been proposed that age-related neural dedifferentiation is not confined to higher-level visual cortex, but also expands to other cortical regions (Koen & Rugg, 2019). For example, in episodic and working memory tasks, multivariate pattern analyses have suggested that age-related activation increases in prefrontal cortex do not contribute additional information to the neural representation (Morcom & Henson, 2018). In general, dedifferentiation arguments attribute areas of increased activation in old age to non-selective neural responses that do not contribute to task processing. This leads to predictions that contrast with the compensatory view. Specifically, if age-related activation of a region is due to dedifferentiated neural responses, its activation strength should not be influenced by task demand and it should always be greater in older people, irrespective of performance differences between age groups.

The compensation and dedifferentiation views have rarely been directly compared. One major reason is that at a population level, healthy individuals exhibit age-related declines in most cognitive abilities, including but not limited to episodic memory, attention and processing speed (Grady, 2012; Salthouse, 2010). As a result, most previous studies were built on cognitive tasks that older adults find more demanding than young people. For these tasks, increased contralateral activation can be interpreted either as compensatory responses to higher task demands or as a breakdown of differentiated neural representations. Therefore, it is critical to test the two competing views with tasks that young people find more difficult than older people, as here the compensation and dedifferentiation accounts predict different results. The compensatory view predicts that the typical pattern of increased contralateral activation in older people will be reversed when young people find a task more challenging than older people. The dedifferentiation view makes no such prediction, since dedifferentiation is not thought to be a product of performance differences.

To adjudicate between the compensation and dedifferentiation accounts, the current study investigated semantic cognition in the ageing brain. Semantic cognition refers to the cognitive processes that store our knowledge about the world (i.e., knowledge representation) and regulate the use of this knowledge to produce task-appropriate behaviours (often termed semantic control). Unlike most cognitive abilities that show a uniform age-related decline, semantic cognition shows a more mixed picture of aging effects. For many years, it has been well-established that the knowledge component of semantic cognition is maintained or even improved with age, as people accumulate knowledge throughout their lives. Older people typically have larger vocabularies and more detailed world knowledge than young people (Kavé & Halamish, 2015; Kavé & Yafé, 2014; D. C. Park et al., 2002; Salthouse, 2004; Verhaeghen, 2003; Wu, Lohani, Homan, Krieger-Redwood, & Hoffman, 2023). In contrast, recent studies have revealed that one aspect of semantic control ability – semantic selection – reduces with age (Hoffman, 2018; Hoffman & MacPherson, 2022; Wu & Hoffman, 2022). Semantic selection refers to the ability to focus on task-relevant aspects of semantic knowledge while inhibiting irrelevant knowledge that inadvertently comes to mind (Badre & Wagner, 2007; Jefferies, 2013). Age-related deterioration in this ability is consistent with, and correlated with, more general declines in executive function and inhibition (Hasher & Zacks, 1988; Hoffman, 2018; Hoffman & MacPherson, 2022). Thus, within the domain of semantic cognition, we were able to test the compensation and dedifferentiation views in a semantic selection task which older people typically find more difficult than young people and in a semantic knowledge task which favours older people.

Laterality effects in semantic cognition are well-characterised, which is another advantage of investigating this cognitive domain. Verbal semantic tasks generate left-dominant activation in a well-established set of semantic regions, often accompanied by weaker recruitment of their right-hemisphere homologues (for meta-analyses, see Hoffman & Morcom, 2018; Jackson, 2021; Rice, Lambon Ralph, et al., 2015). While left hemisphere regions are taken to form the core network for semantic processing, less is known about the functional significance of right hemisphere activation. In the present study, we focus on two major regions involved in verbal semantic processing: the inferior frontal gyrus (IFG) and anterior temporal lobe (ATL) (Jackson, 2021; Rice, Lambon Ralph, et al., 2015).

Left IFG is consistently activated by semantic tasks and is implicated in control processes acting on semantic activation (e.g., semantic selection) (Jackson, 2021; Lambon Ralph, Jefferies, Patterson, & Rogers, 2017). The functional significance of smaller amounts of right IFG activation is a matter of debate. If right IFG contributes to performance in semantic tasks, we would expect activation in this area to increase as a function of task demands. However, fMRI studies in young participants present a mixed picture as to whether right IFG activation does increase as semantic tasks become more difficult (in contrast to left IFG, which reliably shows such effects) (Jung, Rice, & Lambon Ralph, 2021; Krieger-Redwood, Teige, Davey, Hymers, & Jefferies, 2015; Quillen, Yen, & Wilson, 2021). Increased right IFG activation is also frequently reported in people with aphasia following left-hemisphere stroke but the functional significance of this effect is debated. Such effects have been interpreted either as a restoration of language function previously supported by left IFG, or as unhelpful disinhibition of right IFG that makes no functional contribution (Gainotti, 2015; Stefaniak, Halai, & Lambon Ralph, 2020; Turkeltaub, 2015). Finally, healthy older adults tend to show higher right IFG activation than young people during semantic processing (Hoffman & Morcom, 2018). Depending on the functional role of right IFG, this activation could either reflect compensation for reduced efficiency in the core left-lateralised semantic network, in line with the HAROLD view, or a dedifferentiation effect that is not linked to task performance.

The ATL is a key site for semantic knowledge representation (Lambon Ralph et al., 2017). Semantic tasks often recruit the ATLs bilaterally but activation is left-lateralised for single-word comprehension, especially for written words (Hoffman & Lambon Ralph, 2018; Rice, Hoffman, & Lambon Ralph, 2015; Rice, Lambon Ralph, et al., 2015). As with IFG, there is some evidence that right ATL activity may compensate for reduced efficiency in the dominant left ATL. When the function of left ATL is inhibited by transcranial magnetic stimulation, right ATL activation increases, with greater activation predicting better performance (Binney & Lambon Ralph, 2015; Jung & Lambon Ralph, 2016). Increased right ATL activation has also been observed in surgical patients whose left ATL has been removed (Rice, Caswell, Moore, Lambon Ralph, & Hoffman, 2018). In the field of cognitive ageing, very few studies have directly compared differences in ATL activation between age groups, possibly due to the low sensitivity of fMRI to activation in the ventral ATLs (Ojemann et al., 1997). Utilizing source localization techniques, a magnetoencephalography (MEG) study has found that older people recruit the left ATL to a greater extent during semantic processing than younger adults (Lacombe, Jolicoeur, Grimault, Pineault, & Joubert, 2015), which is consistent with a greater reliance on semantic knowledge in older age. Nevertheless, the evidence so far is limited and this finding is restricted to the left ATL, thus it is unknown whether older people show increased right ATL activation, analogous to the shift in laterality seen in the IFGs.

In the present pre-registered fMRI study, we investigated age-related differences in the neural correlates of semantic processing, with a particular focus on right-hemisphere semantic homologues. We used a synonym judgement task that placed high demands on semantic knowledge and a feature-matching task that placed high demands on semantic control. Difficulty in each task was parametrically manipulated, which allowed us to identify task-demand-related activity changes in each age group, both across tasks and across difficulty levels within each task. If the compensation view was correct and the engagement of right IFG was a compensatory response to increased task demands, its activation would increase with difficulty in both age groups. Meanwhile, age differences in activation of this region would depend on age differences in task performance: older people would show more bilateral IFG activation when they found the task more difficult than young people (i.e., the semantic control task), and the reverse effect would be found when the task demands favoured older people (i.e., the semantic knowledge task). Alternatively, if the dedifferentiation account was correct and right IFG’s activation was not related to task demands, this activation would not be regulated by difficulty and older adults would always activate the right IFG more than the young people, irrespective of task performance. We tested parallel predictions for the ATLs, as potential age-related laterality shifts in this region have not previously been investigated.

In addition, we took the opportunity to perform exploratory analyses at a network level, to investigate an emerging domain-general cognitive ageing theory - the Default-Executive Coupling Hypothesis of Aging (DECHA; Spreng & Turner, 2019; Turner & Spreng, 2015). DECHA describes a pattern of changing reliance from fluid intelligence (i.e., problem-solving abilities and executive control) to crystallized intelligence (i.e., stores of prior knowledge) as people age. The theory relates this changed cognitive architecture to three shifts in the functional network architecture of the brain, beyond a single region or hemisphere. First, executive control regions in prefrontal cortex become less strongly modulated by task demands in older people (Cappell, Gmeindl, & Reuter-Lorenz, 2010; Turner & Spreng, 2015), suggesting that older people are less able to recruit cognitive control resources to support performance on challenging tasks. Second, the default mode network (DMN, thought to support prior knowledge and experiences) is less suppressed during task performance in older people, and is also less modulated by task (D. C. Park, Polk, Hebrank, & Jenkins, 2010; Sambataro et al., 2010), suggesting that older people are more reliant on their prior knowledge to support cognition. Third, older people have stronger (but less flexible) connectivity between DMN and executive control regions, which is thought to facilitate their use of prior knowledge to achieve goal-directed behaviours (Spreng & Turner, 2019; Turner & Spreng, 2015). We used network analyses to investigate these effects in the domain of semantic cognition. While previous studies have investigated DECHA predictions in tasks requiring creative thought (Adnan, Beaty, Silvia, Spreng, & Turner, 2019) and autobiographical memory (Spreng et al., 2018), none have investigated the effects of varying demands on knowledge representations and control processes within semantic tasks. Thus, we tested whether older adults would show different patterns of activation and connectivity in the MDN and DMN, as predicted by the DECHA account. We contrasted effects in these networks with the left-hemisphere network supporting semantic control.

## Materials and Methods

Our sample size, hypothesis, study design and analyses were preregistered (available at https://osf.io/pcvfg). The study data are also publicly available (https://doi.org/10.7488/ds/3845 and https://doi.org/10.7488/ds/3846).

### Participants

Forty-five older adults and forty-five young adults were recruited from the Psychology department’s volunteer panel and local advertising, and participated in the study for payment. All participants were native speakers of English and reported to be in good health with no history of neurological or psychiatric illness. As a general cognitive screen, the older participants completed the Mini-Addenbrooke’s Cognitive Examination (M-ACE; Hsieh et al., 2015) prior to starting the experiment. Two older participants scored < 26 of 30 on the M-ACE, and their data were excluded based on our preregistered exclusion criteria. Two young participants’ data were also excluded because of technical issues or structural abnormalities. In the end, data from forty-three older participants (28 females, 15 males; mean age = 68.14 years, *s.d.* = 5.21 years, range = 60 - 79) and forty-three young participants (31 females, 12 males; mean age = 23.07 years, *s.d.* = 3.23 years, range = 18 - 32) were used in the analyses. Levels of formal education were high in both age groups (older adults: mean = 15.65 years, *s.d.* = 2.84 years, range = 10 - 22; young adults: mean = 17.07 years, *s.d.* = 2.53 years, range = 12 - 23), and young adults had completed more years of education than older adults (*t*_84_ = 2.44, two-tailed *p* < 0.05). This reflects greater access to higher education in younger generations. Informed consent was obtained from all participants and the research was performed in accordance with all relevant guidelines/regulations. The study was approved by the University of Edinburgh Psychology Research Ethics Committee.

### Materials

Participants completed three tasks: a semantic knowledge test, a semantic control test and a cognitively demanding non-semantic test (see Figure 1 for examples). The stimuli for all tests were taken from the norms of Wu and Hoffman (2022). The norms define five levels of difficulty (1-5, five for the most difficult) for the stimuli in each test, based on the accuracy and reaction times of participants. By including stimuli with varied difficulty levels, we were able to investigate how brain function changed as a function of demand in different tasks.

**Figure 1.**
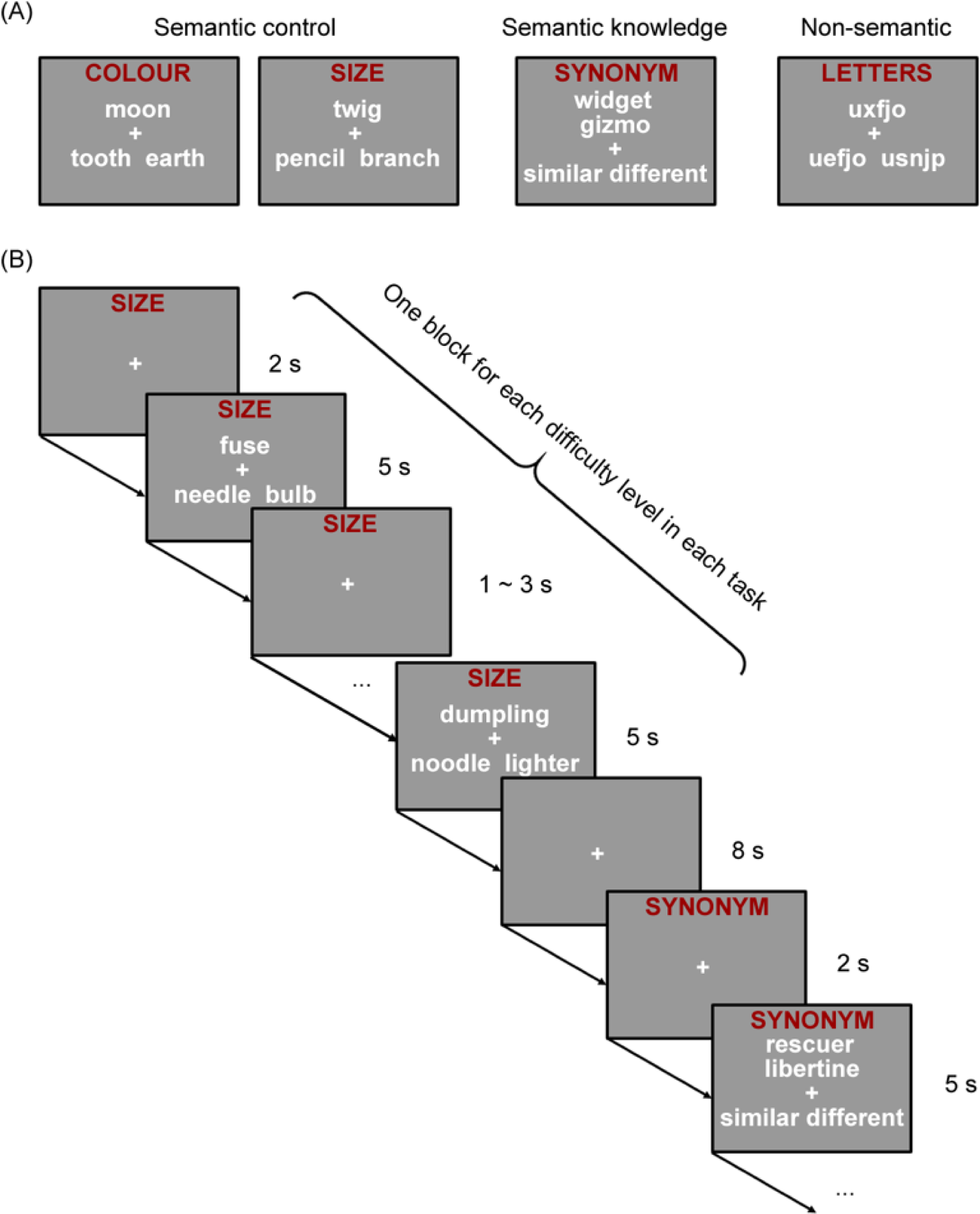
(A) Example items from each task and (B) an illustration of block structure in the experiment.

#### Test of semantic control

Participants completed an 80-item feature-matching task designed to probe their semantic control ability. On each trial of the task, participants were presented with a probe word appearing above the centre of the screen with the two option words in a line below. They were asked to select the option that matched on particular features with the probe (colour on 40 trials and size on 40 trials) from two alternatives. For example, on a colour trial, *moon* would match with *tooth* as both are typically white. This task requires participants to engage semantic control processes, because they must direct their attention to the target semantic properties and inhibit knowledge of other properties, as well as competing but irrelevant semantic associations (*moon-earth*) (Badre, Poldrack, Pare-Blagoev, Insler, & Wagner, 2005; Thompson-Schill, D’Esposito, Aguirre, & Farah, 1997). More difficult trials on this task tend to feature a weaker semantic relationship between probe and target and stronger association between probe and distractor, thus maximising the need for controlled processing (Wu & Hoffman, 2022). Previous studies have found that older people perform more poorly than young people on this task when the competition from irrelevant associations is strong (Hoffman, 2018; Hoffman & MacPherson, 2022).

#### Test of breadth of semantic knowledge

An 80-item synonym judgement task was used to probe semantic knowledge. On each trial, two words appeared above the centre of the screen vertically with the two options (i.e., similar or different) in a line below. Participants were asked to decide if the two words shared a similar meaning or not. More difficult trials tend to feature low-frequency words that are unknown to some people in the population. Thus, this task placed high demands on the semantic knowledge system by requiring comprehension of low-frequency and poorly-understood concepts (Wu & Hoffman, 2022). Previous studies have found that older people perform better than young people on this task (Hoffman, 2018; Hoffman & MacPherson, 2022; Verhaeghen, 2003).

#### Test of non-semantic cognitive control

As a comparison for the semantic tasks, we used an 80-item non-semantic task examining general executive control ability during orthographic processing. The stimuli in this task were 80 triads that consisted of three meaningless letter strings. The triads were presented in a similar fashion to the feature-matching task. Participants were required to choose the option that shared the most letters in the same order as the probe. More difficult trials featured greater similarity between the probe and distractor and decreased similarity between the probe and target, making the distractor harder to reject.

### Design and procedure

There were two scanning runs in the experiment with a block design. In each run, participants viewed 10 blocks from each of the three tasks, i.e., two blocks for each difficulty level of each task. The structure of blocks is shown in Figure 1. Each block started with a 2-s task cue (i.e., “synonym”, “colour”, “size”, or “letters”), which remained at the top of the screen during the whole block. After that, four 5-s task trials were presented, which came from the same difficulty level of the task. The inter-trial interval was 1-3 s (jittered, 2 s on average). The blocks in each run were separated by 8-s periods of fixation.

Block order in each run was randomized. Trial order within each block was pseudo-randomized to make sure that the position of correct responses in the block (i.e., left or right) were balanced. Each stimulus was presented once within the whole experiment. To counterbalance the potential influences of block/trial order on our data, two sets of experimental programs with different block/trial orders were made and each set was used for half of the participants in each age group.

Participants indicated their choice by pressing buttons with their left and right index fingers. For the feature-matching task, the trials in a block were all based on the same feature (colour or size).

### Behavioural data analyses

We used mixed effects models to predict accuracy and RT at the level of individual trials, with age as a between-subjects predictor and task and difficulty level as within-subject predictors (with difficulty as a continuous predictor). We also included trial position in scanning run and position of correct response on screen in each model as covariates of no interest. All mixed effects models in the paper were constructed and tested using the recommendations of Barr, Levy, Scheepers, and Tily (2013) and continuous predictors were standardized prior to entry in the model. Logistic models were estimated for analyses of accuracy and linear models were specified for analyses of RT and neural measures (described later).

We specified a maximal random effects structure for all models, including random intercepts for participants and items as well as random slopes for all predictors that vary within-participant or within-item. For the logistic models, the statistical significance of effects of interest was assessed with likelihood-ratio tests while for linear models, Satterthwaite’s approximation was used with the lmerTest package (Kuznetsova, Brockhoff, & Christensen, 2017). For models including factors with more than two levels, we report the Chi-square (or F value) for tests of the main effects and their interactions. For models where all factors only have two levels, we report effect size (B) and standard error (s.e.) for each effect. Due to the large number of analyses, we report the full tables of all modelled effects in Supplementary Materials. In the main text, we summarise the key effects that are most relevant to our research questions.

### Image acquisition and processing

Images were acquired on a 3T Siemens Skyra scanner with a 32-channel head coil. To minimize the impact of head movements and signal drop out in the ventral temporal regions (Kundu et al., 2017), the study employed a whole-brain multi-echo acquisition protocol, in which data were simultaneously acquired at 3 TEs. Data from the three echo series were weighted and combined, and the resulting time-series were denoised using independent components analysis (ICA). For the functional images, the multi-echo EPI sequence included 46 slices covering the whole brain with echo time (TE) at 13 ms, 31 ms and 50 ms, repetition time (TR) = 1.7 s, flip angle = 73°, 80 × 80 matrix, reconstructed in-plane resolution = 3 mm × 3 mm, slice thickness = 3.0 mm and multiband factor = 2. Two runs of 642 volumes were acquired. A high-resolution T1-weighted structural image was also acquired for each participant using an MP-RAGE sequence with 1 mm isotropic voxels, TR = 2.5 s, TE = 4.4 ms.

Images were pre-processed and analysed using SPM12 and the TE-Dependent Analysis Toolbox (Tedana) (DuPre et al., 2021). The first 4 volumes of each run were discarded. Estimates of head motion were obtained using the first BOLD echo series. Slice-timing correction was carried out and images were then realigned using the previously obtained motion estimates. Tedana was used to combine the three echo series into a single-time series and to divide the data into components classified as either BOLD-signal or noise-related based on their patterns of signal decay over increasing TEs (Kundu et al., 2017). Components classified as noise were discarded. After that, images were unwarped with a B0 fieldmap to correct for irregularities in the scanner’s magnetic field. Finally, functional images were spatially normalised to MNI space using SPM’s DARTEL tool (Ashburner, 2007) and were smoothed with a kernel of 8 mm FWHM.

Data in our study were treated with a high-pass filter with a cut-off of 180 s and the two experimental runs were analysed using a single general linear model (GLM) for each subject. In the GLM, there were regressors for the 3 different tasks (i.e., semantic knowledge task, semantic control task and non-semantic task). For each of these main task regressors, 4 parametric modulators representing the difficulty levels were included. We used simple coding for the modulators of difficulty. By doing this, we captured the general effects of task (averaged across difficulty) with the main task regressors, while the 4 parametric modulators compared activation for difficulty levels 2 to 5 with difficulty level 1. Our pre-registered analysis plan included a regressor coding for attentional lapses during the scans (i.e., failing to respond for 10 or more consecutive trials) but no participant showed such lapses. Each trial was modelled as a separate event of duration 5 s, convolved with the canonical hemodynamic response function. Twelve nuisance regressors modelling movement artifacts, using the three translations and three rotations estimated during spatial realignment, and their scan-to-scan differences, were also included in the GLM.

Contrast analyses were conducted to measure task and difficulty effects at whole-brain level. For each participant, three sets of contrasts that involved the main task regressors were computed: (1) We contrasted each task against fixation/rest; (2) we contrasted each pair of tasks against each other; (3) we contrasted the average of the two semantic tasks with the non-semantic task (for voxel selection in ROI analyses). A set of contrasts that measured difficulty effects were also performed. In these, the modulators for each task regressor were separately compared with fixation to estimate activation for difficulty levels 2 to 5, relative to the baseline of difficulty level 1 in each task. Our pre-registered plan was to first use analysis of variance (ANOVA) to test for differences between difficulty levels 2 to 5, without assuming a linear relationship between activation and difficulty, and then to estimate the linear effects of difficulty on activation. However, our initial analyses indicated that where difficulty effects occurred, they were always approximately linear in nature, hence we only report effects of the linear contrast here.

### Definition of pre-registered regions of interest (ROIs)

To analyse effects in ATLs and IFGs, we used a method that combined anatomical masks with selection of task-activated voxels at an individual subject level. Several steps were involved in this procedure.

First, anatomical ATL and IFG masks were defined (Figure 4A). We defined IFG with the Harvard - Oxford structural atlas (http://fsl.fmrib.ox.ac.uk/fsl/fslwiki/Atlases). We included BA 44/45/47 (pars opercularis, pars triangularis and pars orbitalis), limiting BA47 to voxels more lateral than x = +/− 30 to restrict this large area to its semantic-related parts (Asyraff, Lemarchand, Tamm, & Hoffman, 2021; Hoffman, 2019; Hoffman & Tamm, 2020). We defined ATL in a similar fashion to a previous study (Hoffman & Lambon Ralph, 2018). We first generated masks of the inferior temporal gyrus, fusiform gyrus, superior temporal gyrus, and middle temporal gyrus by including all voxels with a greater than 50% probability of falling within the corresponding areas in the LONI Probabilistic Brain Atlas (LPBA40). Then we divided each of these masks into 6 roughly-equal-length sections that ran along an anterior-to-posterior axis (section 0-5, 0 for the most anterior), with divisions approximately perpendicular to the long axis of the temporal lobe. Lastly, we constructed left and right ATL ROIs by combining sections 1 and 2 of the above masks.

Second, within the pre-defined anatomical ATLs and IFGs, for each participant, we used the contrast of the semantic knowledge and control tasks (average) minus the non-semantic task to identify voxels that were most responsive to semantic processing (relative to domain-general cognitive demands). Within each anatomical ROI, we selected the top 20% most active voxels for semantic processing for each individual participant and used these voxels as our final ROI to extract effects from. This means that analyses of IFG and ATL focus on those parts that were most selective for semantic processing in each participant.

### ROI-level analyses

We extracted voxel-average activation estimates for the task and difficulty regressors in the pre-registered ROIs defined for each participant. We then conducted a series of mixed model analyses to investigate effects of *age, task* and *difficulty* in the ROIs. In our pre-registration, we planned to use ANOVA to perform these analyses, treating difficulty as an unordered factor with four levels. However, effects of difficulty were largely linear in form so we decided that a linear mixed effects approach was better suited to modelling these effects, as difficulty could be included as a continuous predictor. Thus, we report results of mixed effects models here (ANOVAs produced qualitatively similar results). For the task-based (but not difficulty-based) models, we were only able to include random intercepts for participants due to the restricted number of observations.

### Whole-brain-level analyses

In this section of analyses, we submitted the first-level whole-brain activation results to second-level analyses for group effects. First, by conducting one-sample t-tests over the maps that contrasted each of the tasks with fixation and with each other, we generated task-related activation maps for each task within each age group. Second, two-sample t-tests were computed to examine the activation differences between older and young people during semantic processing, for the semantic control minus non-semantic and semantic knowledge minus non-semantic contrasts. Finally, similar to our ROI-based analyses, we investigated the linear effects of difficulty for each task and each age group. One-sample t-tests were computed over the linear contrast maps of the 4 difficulty modulators in each task to test for effects within each age group. Two-sample t-tests were used to test age-related differences in difficulty effects. Whole-brain effect maps were thresholded at a voxelwise p < 0.005 and corrected for multiple comparisons at the cluster level using SPM’s random field theory (FWE-corrected p < 0.05).

### Network-level exploratory analyses

#### Definition of network masks

We defined 3 brain networks for our exploratory analyses. The multiple-demand network (MDN) was obtained from Fedorenko, Duncan, and Kanwisher (2013), and consisted of voxels showing a positive response to cognitive demands in multiple domains. The semantic control network (SCN) was obtained from the meta-analysis of Jackson (2021). This meta-analysis identified regions showing responses to cognitive control demands in semantic tasks. Last, we defined the default mode network (DMN) using data from Yeo et al. (2011), which identified this network based on parcellation of resting-state fMRI connectivity data. A small proportion of voxels were shared across the three networks. To ensure independence of the networks, voxels that fell within the SCN were excluded from the DMN and MDN, and voxels that fell within the MDN were then excluded from the DMN. The maps of the three networks of interest are shown in Figure 8A.

#### Network timecourse extraction

There were two parts to the network-level analyses. In the first part, we examined activation effects in the networks of interest as a function of age, task and difficulty. In the second part, we investigated age-related changes in network connectivity for semantic relative to non-semantic processing. Analyses were performed on BOLD timecourses of the three networks, which were extracted with the spatially-constrained ICA function in the GIFT toolbox (https://trendscenter.org/software/gift/). ICA assumes that fMRI signals consist of a mixture of spatially (or temporally) independent components, and the aim of ICA is to recover those components (Calhoun, Liu, & Adali, 2009). Spatially-constrained ICA is a semi-blind ICA algorithm in which the components are extracted under the constraint that they conform to specific spatial templates (Lin, Liu, Zheng, Liang, & Calhoun, 2010). Using our network masks (i.e., the MDN, SCN and DMN) as the spatial templates, we performed spatially-constrained ICA on each participant’s data. This method gave us three independent timecourses from each participant’s data, corresponding to the three networks. Further analyses were conducted on these extracted component timecourses.

#### Network-level activation

For each participant, we built a GLM with the same structure as earlier analyses to model the task regressors and difficulty modulators. However, here we used the GIFT toolbox to fit the model to the extracted ICA timecourse for each network, instead of the timecourses from individual voxels. Therefore, the parameter estimates for the task regressors and difficulty modulators in this model represent the influences of task and difficulty on network activations. We used a series of mixed effects models to identify the significant *task-based* and *difficulty-based* network activation effects at the group level.

#### Network-level functional connectivity

Here, we investigated how semantic processing affected connectivity between the three networks of interest, relative to non-semantic processing. We used generalised psychophysiological interactions (gPPI) to achieve this objective (McLaren, Ries, Xu, & Johnson, 2012). PPI measures how the intrinsic correlation between two brain regions (i.e., the physiological effect) changes when a participant engages in different cognitive activities (i.e., the psychophysiological interaction). While traditional PPI can only assess connectivity differences between two task conditions, gPPI allows for the inclusion of more than two task conditions in a single PPI model, as is required in the current study. We used the gPPI toolbox (https://www.nitrc.org/projects/gppi) to perform these analyses.

We first built a simplified GLM for each participant, which included the task regressors but not the difficulty modulators, and used the gPPI toolbox to add the seed timecourse and its interaction with the task regressors (the PPI terms). The timecourse for the seed was the network component timecourse extracted with GIFT (for similar approaches, see Goulden et al., 2014; Lee Masson, Op de Beeck, & Boets, 2020). Therefore, the PPI terms estimated how the connectivity between the seed network and each voxel in the brain changed for each task relative to fixation. By contrasting the model estimates, we were able to test how connectivity varied with task (e.g., semantic control_PPI_ minus non-semantic_PPI_). Importantly, because we were interested in the functional interactions between networks, we used the GIFT toolbox to fit the above GLM to the component timecourse of a target network instead of the timecourse from a single voxel. For instance, a model with the DMN as the seed could be used to predict the timecourses of the SCN or MDN, and the obtained model estimates indicated how the correlation between DMN and SCN (or MDN) was affected by the experimental tasks. As PPI effects are not directional (O’Reilly, Woolrich, Behrens, Smith, & Johansen-Berg, 2012), we averaged the effects across models for each pair of networks so that each network in the pair had the same chance serving as the seed and target.

After constructing the network gPPI models and obtaining the model estimates, we then used mixed effects models to investigate how functional connectivity between each pair of networks was influenced by semantic control and knowledge processing for different age groups.

## Results

### Pre-registered analyses

#### Behavioural data

We used a series of mixed effects models to investigate participants’ performance on the three tasks in the scanner (Supplementary Table S1). Estimated effects from the models for individual tasks are shown in Figure 2. Older people were slower to respond than young people in all tasks (all *p* < 10^−4^), indicating an age-related decline in general processing speed. Age × task interactions for both accuracy (*p* < 0.05) and RT (*p* < 10^−5^) indicated that age had differing effects on the three tasks. In accuracy, as expected, older people outperformed young people on the semantic knowledge task (*p* < 0.01). There were no age group differences in the semantic control task (*p* = 0.708) or the non-semantic task (*p* = 0.426). Age effects on RT were also smallest in the semantic knowledge task (age x task interaction, *p* < 10^−5^). This suggests that older people found the semantic knowledge task less demanding than young people, while this was not true for the other tasks.

**Figure 2.**
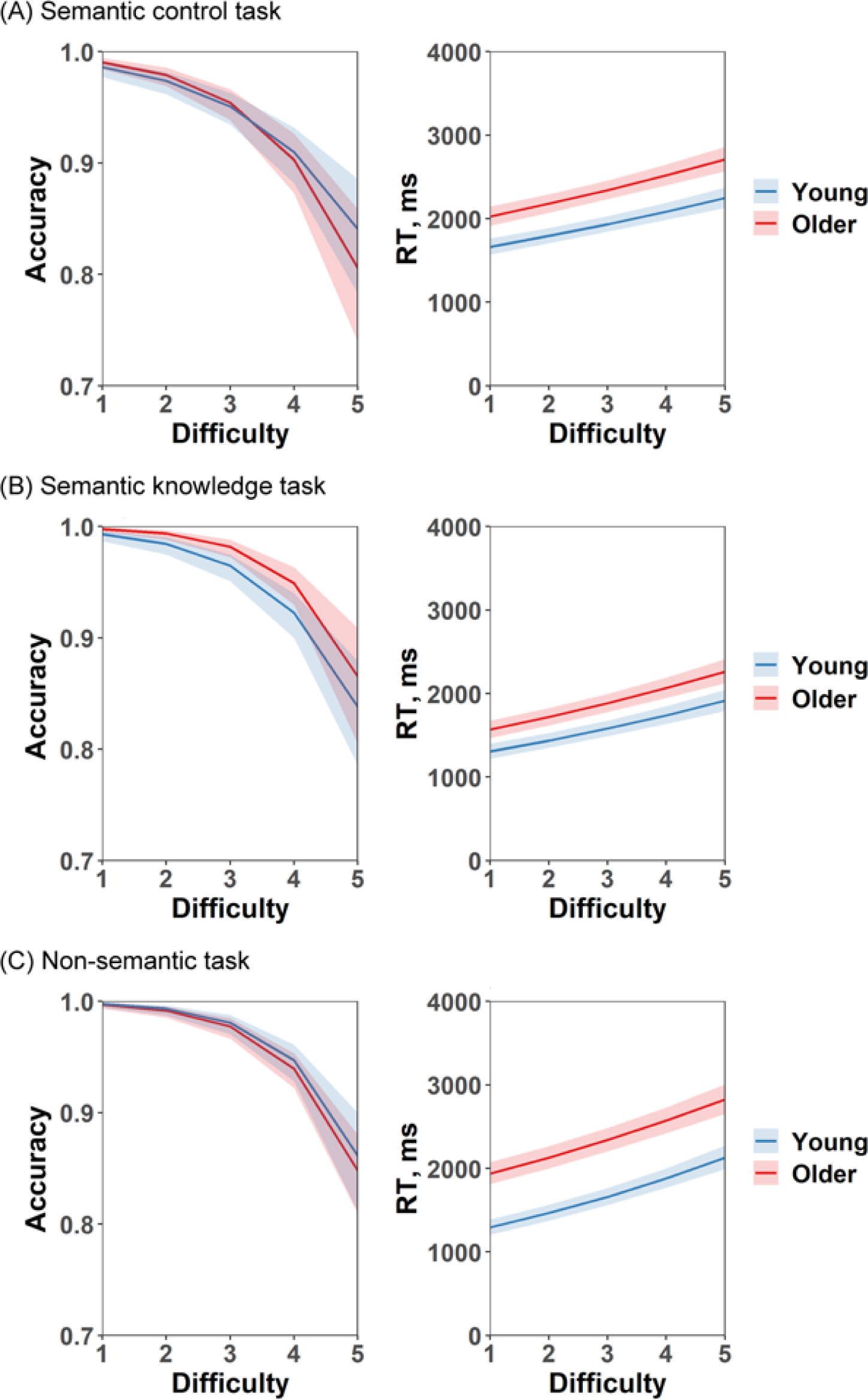
Modelled effects of age and difficulty on accuracy and RT in each task. Shadow areas indicate 95% confidence intervals.

The difficulty manipulations were successful, having significant effects on accuracy and RT in all tasks (all *p* < 10^−15^). The only interaction between difficulty and age was for RT in the non-semantic task (*p* < 0.001), where difficulty had a stronger influence on RTs in young people. For the semantic tasks, the difficulty manipulations had similar effects on both age groups.

### ROI-level analyses

#### Semantic processing in bilateral ATL and IFG

Before examining the compensation and dedifferentiation hypotheses in our ROIs, we first conducted a set of analyses to test the roles of ATL and IFG in different semantic tasks and in different age groups.

As outlined in the Introduction, models of semantics link the representation of semantic knowledge with ATL and semantic control processing with IFG (Lambon Ralph et al., 2017; Patterson, Nestor, & Rogers, 2007). Therefore, we predicted stronger activation in ATL for the knowledge task cf. the semantic control task and the reverse in IFG. Previous studies have also suggested that there is a shift toward greater reliance on knowledge and less on executive control in older adulthood (Spreng & Turner, 2019; Turner & Spreng, 2015). Thus, it is possible that older people would rely more on the ATL than IFG in comparison with the young. To test these two predictions, we fitted a mixed effects model predicting activation effects from age (young vs. older), task (knowledge vs. control) and ROI (ATL vs. IFG, collapsed over hemispheres). Consistent with the first prediction, ATL was more engaged in the knowledge task and IFG was more activated in the semantic control task (ROI × task effect, *p* < 10^−4^, see Supplementary Table S2 and Figure 3). However, our dataset revealed no age differences in the activation in the semantic ROIs (age main effect, *p* = 0.443; age × ROI, *p* = 0.066).

**Figure 3.**
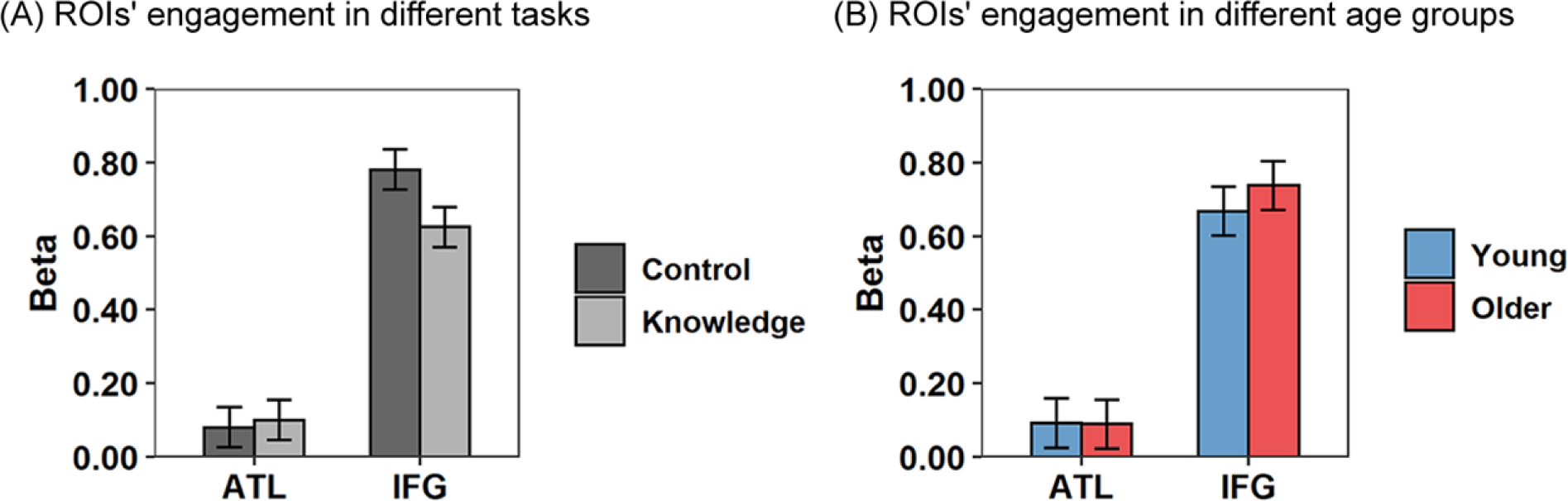
An investigation of shifting division of labour between the IFG and ATL in the semantic tasks. We tested (A) modelled effects of task and ROI on activation (vs. fixation), and (B) modelled effects of age and ROI on activation (vs. fixation). Error bars indicate 95% confidence intervals.

The above results suggest a division of the roles of ATL and IFG in semantic tasks, in which IFG engaged more in the semantic control processing and ATL more in knowledge. However, the overall age-related difference in activation in the ATL and IFG was not significant in our dataset, indicating that a shift towards reliance on prior knowledge in older people might be supported by a wider range of regions (i.e., networks) rather than a single area (e.g., ATL). Furthermore, as these analyses averaged across hemispheres, they did not test for age-related changes in laterality. The next set of analyses addressed this question.

#### Test of the compensation and dedifferentiation hypotheses

In this section, we investigated the two main accounts with a series of linear mixed effects models built to probe *task-based* and *difficulty-based* effects in the left and right IFG and ATL (Figure 4A). For the task-based analysis, the compensation hypothesis predicted that young people would show more right-hemisphere activation than older people for the knowledge task (as young people found this task more difficult) while the reverse would be true for the control task. The dedifferentiation view predicted increased right-hemisphere activation in older people irrespective of task. To test these predictions, we fitted linear mixed effects models with age, task and hemisphere as predictors to predict the activation effects in the semantic tasks. The results are shown in Supplementary Table S3 and Figure 4B. Contrary to the predictions of both theories, we found no age differences in overall activation (age main effects, *p*_IFG_ = 0.269, *p*_ATL_ = 0.950) and no age × hemisphere interactions (*p*_IFG_ = 0.290, *p*_ATL_ = 0.148), indicating that neither the magnitude nor laterality of activity in these regions was affected by age. This suggests that age-related changes in activation in our study, if present at all, occurred outside the core regions for semantic processing.

**Figure 4.**
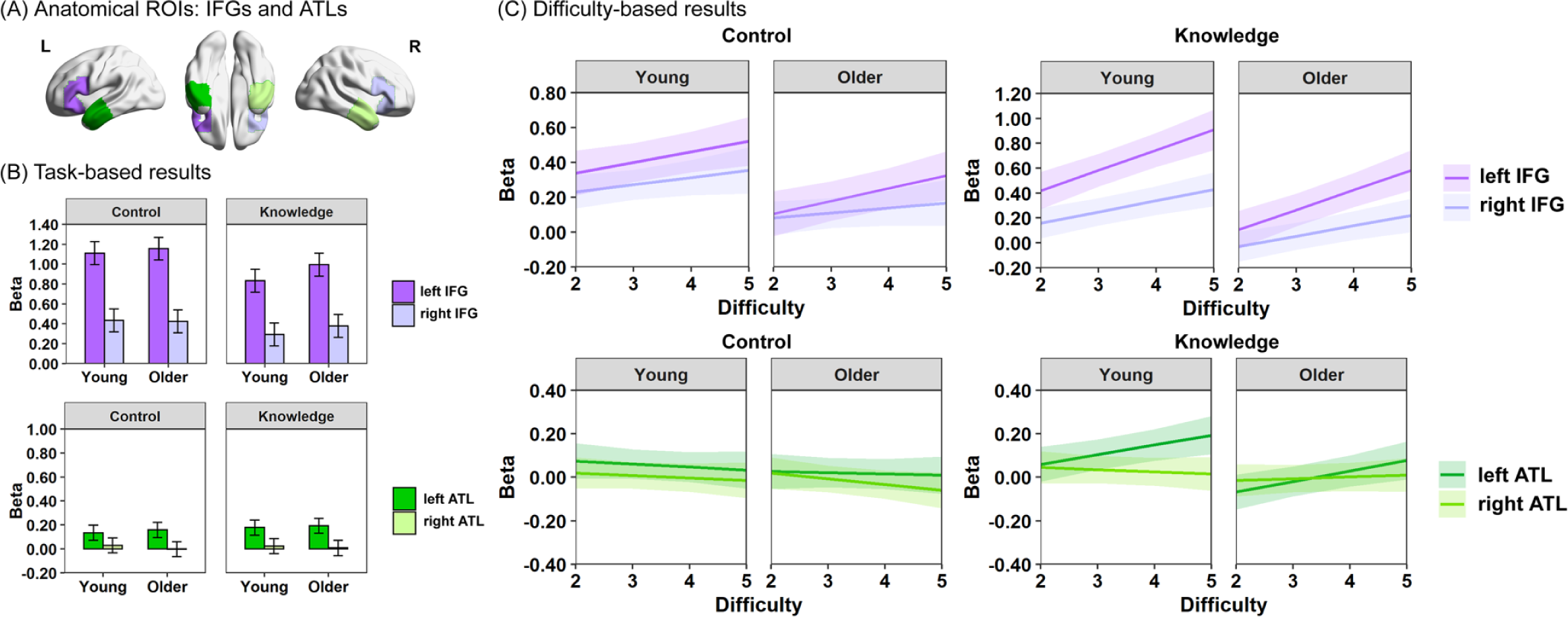
Results in regions of interest. (A) Anatomical ROIs were specified for left and right IFGs (dark and light purple) and left and right ATLs (dark and light green). (B) Modelled effects of age, task and hemisphere on activation (vs. fixation) in the IFGs and ATLs separately. Error bars indicate 95% confidence intervals. (C) Modelled effects of age, difficulty and hemisphere on activation (vs. difficulty level 1) in the IFGs and ATLs and for the semantic control task and semantic knowledge task separately. Shadow areas indicate 95% confidence intervals.

In the IFGs, both left and right regions were activated above baseline (fixation) but with a pronounced left-hemisphere bias (hemisphere main effect, *p* < 10^−15^). There was more activation for the semantic control task (task main effect, *p* < 10^−6^) and this task also showed a stronger left-hemisphere bias than the knowledge task (hemisphere × task effect, *p* < 0.05). The ATL also showed a left-hemisphere bias (*p* < 10^−15^) but this did not depend on task (*p* = 0.203). Right ATL was not activated above baseline in either task; thus, only the left ATL appeared to contribute to performance on these semantic tasks.

For the difficulty-based analysis, the compensation theory predicted that activation in the right IFG/ATL of both age groups would increase with task demands, in a similar fashion to their left hemisphere homologues. However, if right-hemisphere over-activation in older people is due to dedifferentiation of neural responses, these regions would not show increased activations with task difficulty, as they would not contribute to task performance. To test these hypotheses, we fitted a mixed effects model for each pair of ROIs and each semantic task, which used age, difficulty and hemisphere as predictors. As shown in Supplementary Table S4 and Figure 4C, the IFGs increased their activation with difficulty in both semantic control and knowledge tasks (difficulty main effects, both *p* < 0.001), but the ATL only showed a significant difficulty effect for the knowledge task (*p*_control_ = 0.147, *p*_knowledge_ < 0.05). In the knowledge task, both IFG and ATL exhibited a stronger difficulty effect in the left hemisphere than the right (hemisphere × difficulty effect, both *p* < 0.001).

In a post hoc analysis, we tested difficulty effects for the left and right IFG/ATL separately. In the IFGs, both left and right hemispheres showed significant increases with difficulty in both tasks (all *p* < 0.05). In the ATLs, however, a significant effect of difficulty was found in the left ATL for the knowledge task (*p* < 10^−4^) but not the control task (*p* = 0.393). Activation in the right ATL was not influenced by difficulty in either task (*p*_knowledge_ = 0.919, *p*_control_ = 0.054 but in opposite direction to prediction). These results support the compensation view of right IFG activation, since this region showed demand-related increases in both tasks. The picture in ATL was more complex, with only the left hemisphere showing a demand-related response, and only then in the knowledge task. This suggests that, in contrast to IFG, right ATL is not recruited to support performance in demanding semantic tasks of the kind used here.

Finally, we did not find age-related differences in the effects of difficulty across ROIs and semantic tasks (all main and post hoc models, age × difficulty effect, all *p* >= 0.326), even though young people’s IFGs exhibited a greater increase in activation overall relative to the lowest difficulty level (all main and post hoc models, age main effect, all *p* < 0.05).

#### Test of the relationship between behavioural performance and task activation

In the last section of the ROI-level analyses, we explored the relationship between individual behavioural performance and ROI activation. To do this, we computed a performance score for each participant that reflected their accuracy and speed on each task. We began by calculating their mean accuracy and RT (on correct trials) across all items in the task. Then for each task, we z-transformed the individual accuracy and RT across all participants in the same age group and subtracted the Z-score of RT from the Z-score of accuracy for each subject as the performance measure. Thus, participants who had relatively high accuracy rates and relatively low RTs received higher performance scores. Then correlations between the performance score and ROI activation were calculated for each task and each age group. The results are reported in Supplementary Table S5. We found that the activation in the left ATL was significantly positively correlated with the behavioural performance of the older participants in the semantic control task and non-semantic task (both *p* < 0.05). However, there was no other significant effects for the ATLs or IFGs in either age group (all *p* >= 0.09).

#### Summary

Our pre-registered ROI-level results showed that both left and right IFGs responded to changed semantic demands across age groups and tasks. This supports the compensation view of right IFG activity and suggests that in the domain of semantic cognition, activation increases in right IFG in older people cannot be simply interpreted as maladaptive dedifferentiation. However, the age-related laterality differences predicted by the compensation account were not found. This suggests that age-related changes in this dataset occurred outside core semantic processing regions. Therefore, analyses to investigate functional reorganization at a larger scale (i.e., whole-brain and network levels) were conducted in the following sections.

### Whole-brain-level analyses

Figure 5 shows activation for each semantic task versus the non-semantic task. The two age groups showed largely similar patterns. For the semantic control versus non-semantic contrast, both age groups showed positive effects in semantic regions including IFG, ATL, posterior lateral temporal lobe and angular gyrus, and older people also showed activation in parts of the DMN such as the posterior cingulate and ventromedial prefrontal cortex. In contrast, higher activation for the non-semantic task was found for both age groups in large areas of parietal and frontal lobes, partly overlapping the MDN. The semantic knowledge versus non-semantic contrast revealed similar sets of regions, but the IFGs’ preference for semantic processing became less prominent here, mirroring the ROI results and reflecting less emphasis on semantic control ability in the knowledge task. Contrasts of knowledge versus control and of each task versus fixation are presented in Supplementary Figure S1.

**Figure 5.**
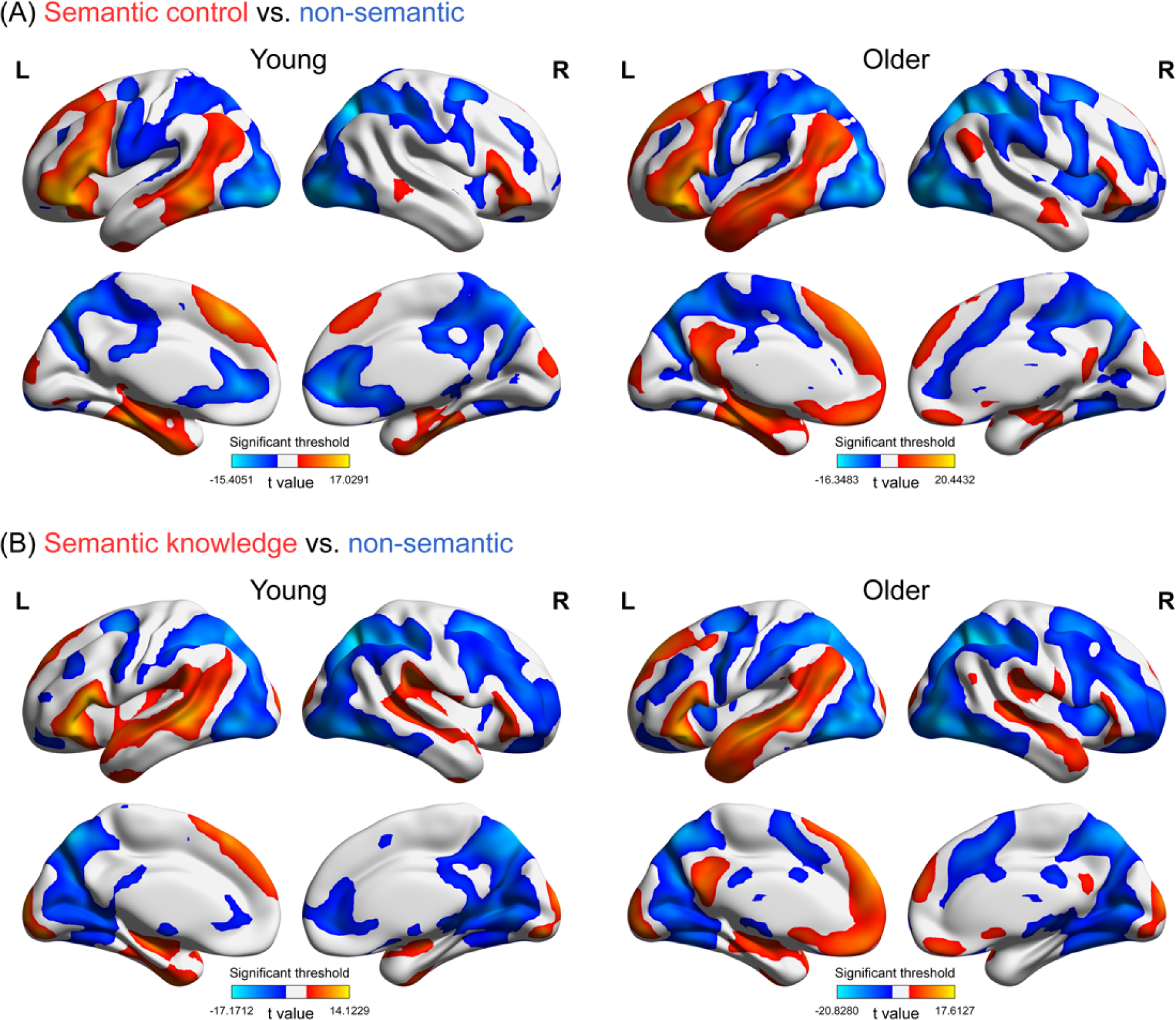
Results of whole-brain univariate activation analysis (task effects, one-sample t test). This figure shows the univariate group activation maps for the contrasts of (A) semantic control task versus non-semantic task and (B) semantic knowledge task versus non-semantic task. Results were corrected for multiple comparisons, voxelwise p < 0.005, FWE corrected cluster threshold p = 0.05.

Next, to evaluate age-related differences, two-sample t tests were computed over the contrast maps of semantic control versus non-semantic and semantic knowledge versus non-semantic. There was a similar pattern of age effects in the two contrasts (Figure 6). Older people activated medial parts of the DMN (posterior cingulate and ventromedial prefrontal cortex) more than young people when making semantic judgements. They also showed more activation in some lateral parts of ATL. In contrast, young people activated areas such as the insulae, pre-supplementary motor area and posterior inferior temporal gyrus more than older people. These regions are parts of the MDN, a set of executive function regions that respond to increased cognitive demands across a range of domains (Duncan, 2010; Fedorenko et al., 2013).

**Figure 6.**
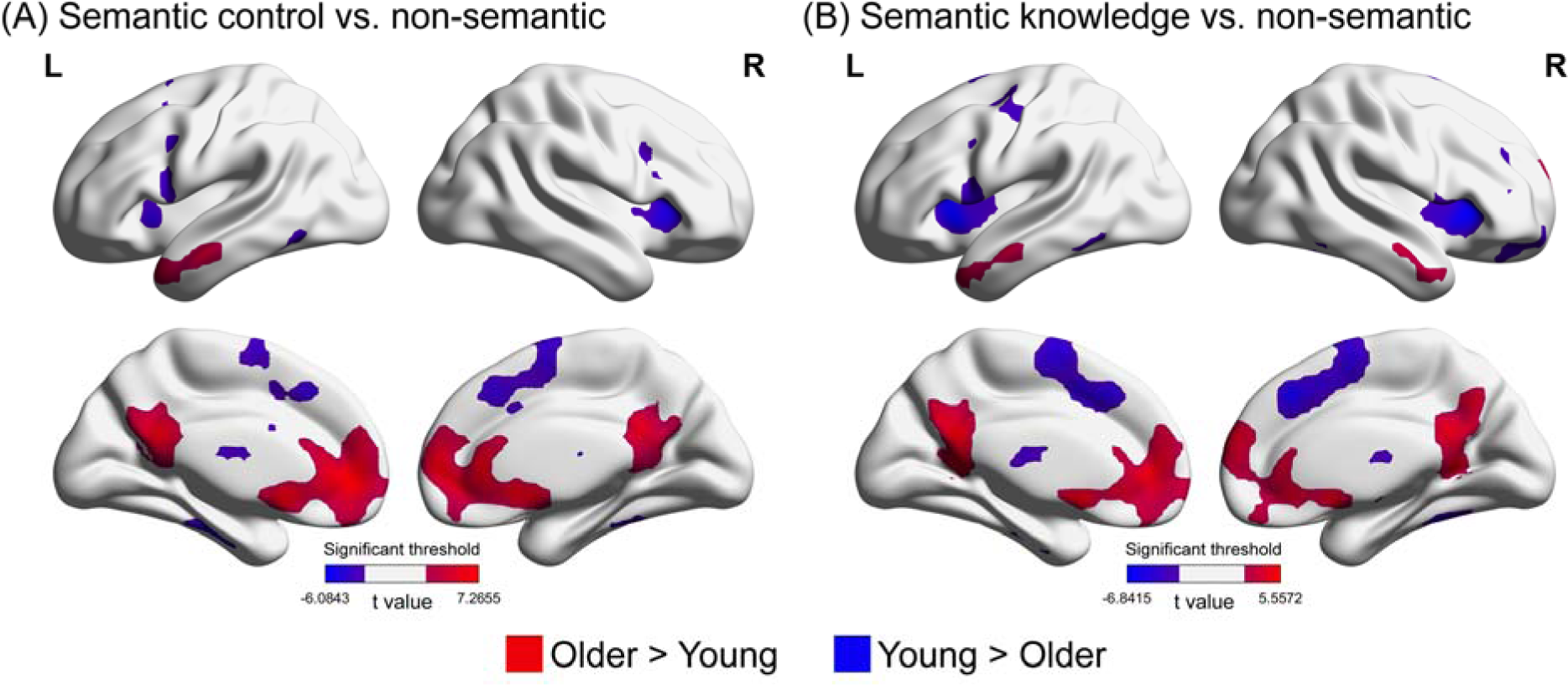
Effects of age on semantic activations. This figure shows the univariate age group activation comparison (older vs. young) for the contrasts of (A) semantic control task versus non-semantic task and (B) semantic knowledge task versus non-semantic task. Results were corrected for multiple comparisons, voxelwise p < 0.005, FWE corrected cluster threshold p = 0.05.

Finally, we investigated task difficulty effects in each task and age group. Figure 7 shows the results, with hot colours indicating regions whose activation increased with difficulty and cold colours showing decreases. For the semantic control task, both age groups showed increasing activation in semantic control and domain-general control areas as difficulty increased, including lateral prefrontal regions, posterior inferior temporal gyrus, insulae, intraparietal sulcus and pre-supplementary motor area. There were no age-related differences when we compared across age groups. For the semantic knowledge task, similar regions showed increased activation as difficulty increased, but DMN areas such as the angular gyrus, posterior cingulate and ventromedial prefrontal cortex showed strong deactivation effects in the young group. A comparison of age groups demonstrated that these deactivation effects were significantly smaller in older adults. For the non-semantic task, in both age groups a wide range of executive control areas across the frontal, parietal and occipital lobes showed greater activation when the task was more demanding. In contrast, DMN regions including the ATLs, angular gyrus, posterior cingulate and ventromedial prefrontal cortex showed demand-related deactivation, especially in the young group. The group comparison suggested that older people showed less deactivation in some DMN areas (e.g., posterior cingulate and ventromedial prefrontal cortex) as a function of task demands, while MDN regions (posterior inferior temporal and intraparietal sulcus) were more responsive to non-semantic task demands in the young people.

**Figure 7.**
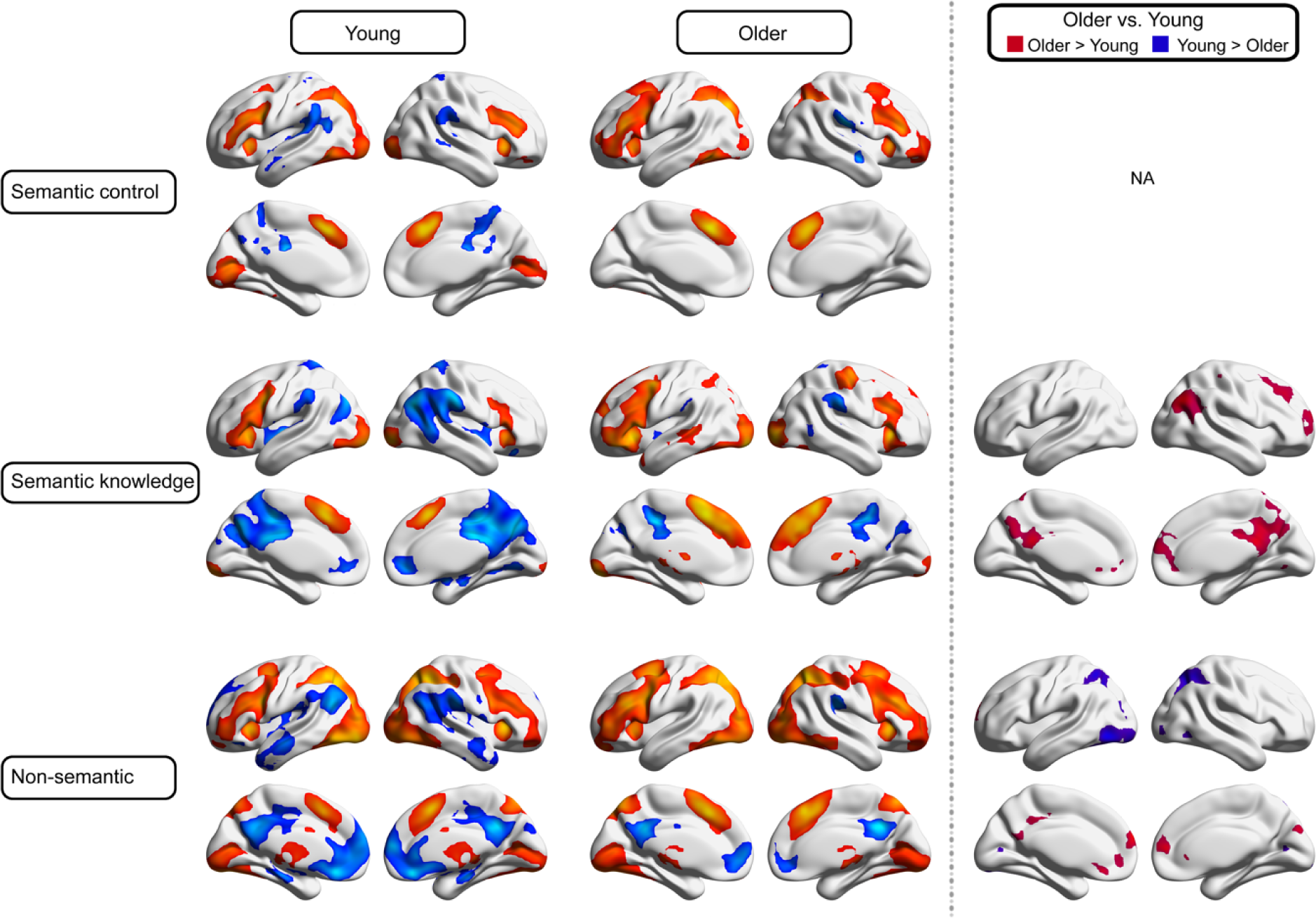
Results of whole-brain univariate activation analysis (difficulty effects, one-sample and two-sample t tests). This figure shows the univariate group activation maps for each age group and the comparison between older and young people, for the linear difficulty effects in each task. In the first two columns, hot colours indicate increases in activation for increasing difficulty and cold colours indicate decreases. Results were corrected for multiple comparisons, voxelwise p < 0.005, FWE corrected cluster threshold p = 0.05.

#### Summary

Although recruitment of ATL and IFG did not differ between age groups, there were clear age differences at the whole-brain level which appeared to map to the two well-studied brain networks (DMN and MDN). To investigate these effects further, we conducted network-level exploratory analyses in the next section.

### Network-level analyses

In this section, we explored the functional reorganization of the ageing brain in three distributed brain networks (shown in Figure 8A). Guided by the whole-brain results, we investigated effects in (1) MDN, which is thought to support task-related executive control processes across multiple domains (Duncan, 2010; Fedorenko et al., 2013) and (2) DMN, which includes parts of the ATLs and is thought to support access to and use of prior knowledge, including episodic and semantic memories (Binder & Desai, 2011; Binder, Desai, Graves, & Conant, 2009; Binder et al., 1999). The DECHA theory proposes that reliance on these systems shifts with age, with older people relying more on access to prior knowledge and less on executive control to complete tasks (Spreng & Turner, 2019; Turner & Spreng, 2015). We also investigated effects in (3) the semantic control network (SCN; Jackson, 2021). This is a set of regions, centred on left IFG, which reliably respond to increased executive demands in semantic tasks. It includes both domain-general and semantic-specific regions (Jackson, 2021). We included it to determine if this semantically-tuned executive network would show similar age-related effects as the domain-general MDN.

**Figure 8.**
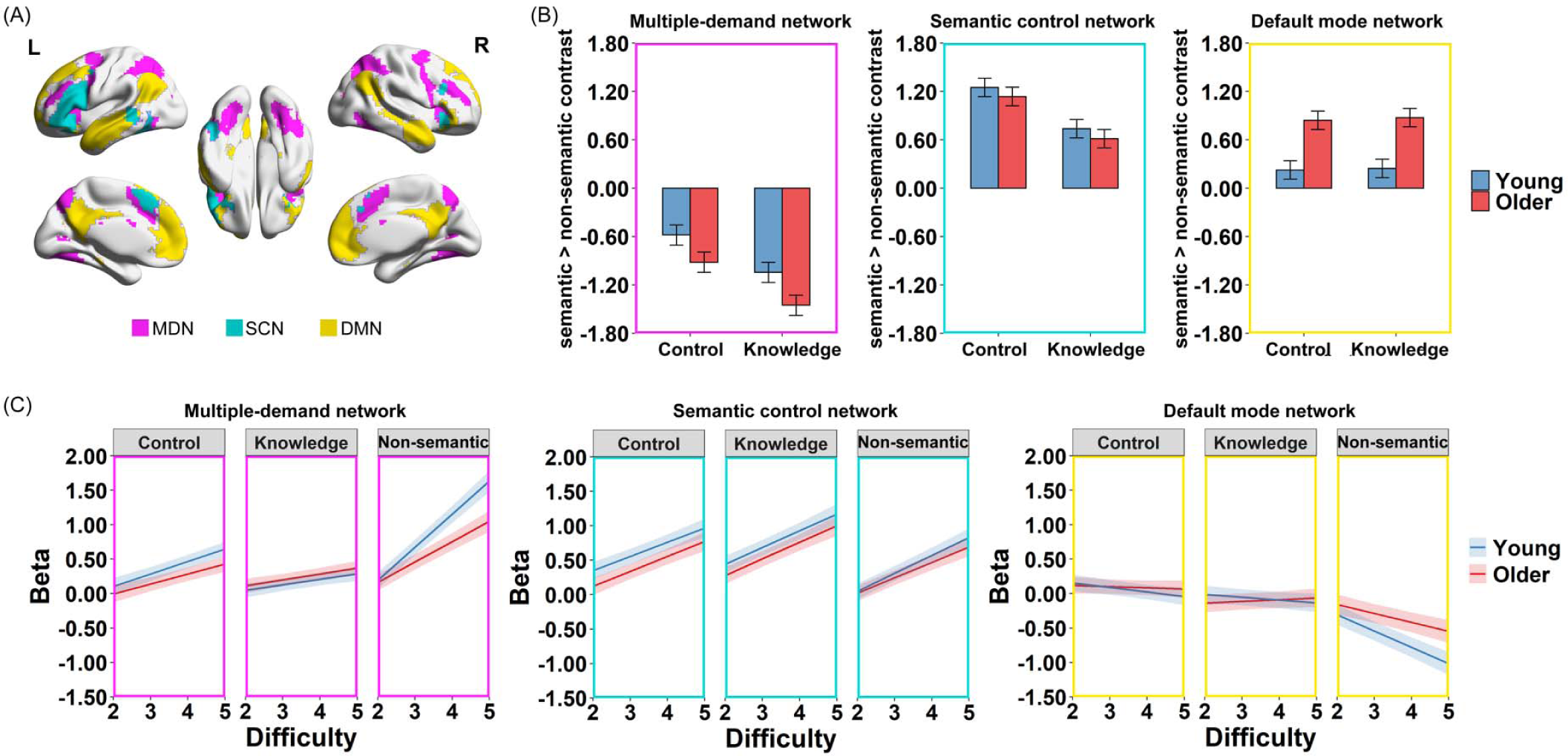
Results in networks of interest. (A) Masks used for the network-level analyses, including MDN (pink), SCN (teal) and DMN (yellow). (B) Modelled effects of age and task on neural activation (vs. the non-semantic task) in each network. Error bars indicate 95% confidence intervals. (C) Modelled effects of age, task and difficulty on neural activation (vs. difficulty level 1) in each network. Shadow areas indicate 95% confidence intervals.

#### Network-level activation

As for our ROI analyses, we first conducted a set of *task-based* analyses, which examined if the two age groups engaged MDN, SCN and DMN differently in different semantic tasks. In each network, we contrasted individual-level beta values for each semantic task with the non-semantic task, and we used the resulting contrast values to fit a mixed effects model with age and task as predictors. The results are shown in Supplementary Table S6 and Figure 8B. MDN was more engaged in semantic control than knowledge processing (*p* < 10^−15^) although it was most responsive to non-semantic processing, reflected by the overall negative activation effects. In addition, this network was more activated for semantics in young people than older people in the semantic tasks (*p* < 10^−4^). This suggests that young people are more likely to recruit domain-general executive resources to support performance in semantic tasks. The SCN was positively engaged by semantic tasks and was more active in the semantic control task than the knowledge task (*p* < 10^−15^). This is consistent with its established role in semantic control functions. Recruitment of SCN for semantic processing did not differ between age groups (*p* = 0.133). Finally, the DMN was also positively engaged by semantic processing, but with no difference between control and knowledge tasks (*p* = 0.300). Older people activated this network more than the young people (*p* < 10^−11^), suggesting that older people were more reliant than young people on the functions of this network when making semantic decisions. No age × task interaction effects were found in any network (*p* >= 0.119).

For *difficulty-based* analyses, we investigated if the activation in each network was influenced by task demands and if these effects differed between age groups. Here, we built mixed models with age, task and difficulty as predictors of activation in each network, with the easiest difficulty level in each task as the baseline. The results are shown in Supplementary Table S7 and Figure 8C. MDN’s activation increased with difficulty across semantic and non-semantic tasks (difficulty main effect, *p* < 10^−15^). However, this network was more sensitive to non-semantic demands than to demands in semantic tasks (task × difficulty effect, *p* < 10^−15^). Previous studies have shown that MDN regions are less modulated by task demands in older people (Cappell et al., 2010; Persson et al., 2004; Turner & Spreng, 2015), suggesting that older people are less able to recruit these regions to support task performance. Here, however, this was only true in the non-semantic task. In contrast, both age groups exhibited similar effects of difficulty in the semantic tasks (age × task × difficulty effect, *p* < 10^−4^).

The SCN also showed a generally positive response to increasing difficulty across tasks (difficulty main effect, *p* < 10^−15^), but here the difficulty effect did not vary with task or age group (all *p* >= 0.286). Lastly, the DMN showed an overall decrease in activity as difficulty increased (difficulty main effect, *p* < 10^−12^), but this effect was largely confined to the non-semantic task (task × difficulty effect, *p* < 10^−12^). Older people’s DMN was less deactivated as a function of difficulty than the young people (age × difficulty effect, *p* < 0.001). This effect was most obvious in the non-semantic task, where activity declined with difficulty in both groups but more strongly in the young people. In the semantic tasks, there was a slight negative effect of difficulty in the young group but either no effect or a slight positive effect in the older group.

#### Network-level functional connectivity

Finally, we explored how the interaction between each pair of networks varied with task and how healthy ageing influenced these between-network interactions. Here, gPPI was used to measure how functional connectivity between networks changed when participants engaged in semantic processing (with the non-semantic task as the baseline). We built a mixed effects model for each pair of networks with age and task as predictors of the PPI connectivity effect. As shown in Supplementary Table S8 and Figure 9, the DMN was more connected to the SCN and MDN during semantic tasks (cf. non-semantic). This indicates that semantic tasks elicited increased interaction between DMN and both semantic and domain-general control networks. In contrast, connectivity between SCN and MDN was unchanged during semantic processing (in the control task) or showed a slight reduction (in the knowledge task). All three networks were more strongly connected with each other in the semantic control task than the knowledge task (task main effect, all *p* < 10^−4^), suggesting that the control task required more interaction between knowledge-supporting and executive control regions. There were no overall age differences in inter-network connectivity (age main effect, all *p* >= 0.077). There was, however, an age × task interaction in MDN-SCN coupling (*p* < 0.05). During the knowledge task, these networks decoupled more in the young group. This might indicate a greater functional separation in the operation of domain-general and semantic control systems in young people.

**Figure 9.**
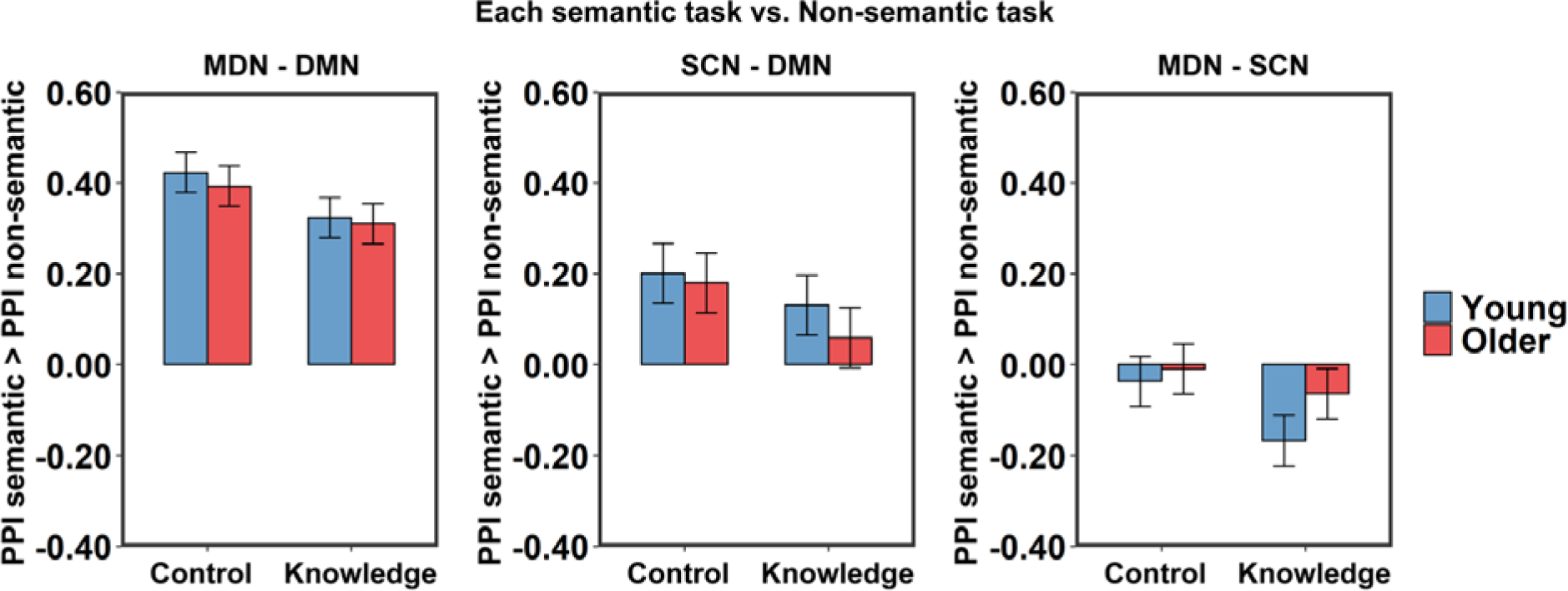
Modelled effects of age and task on the PPI functional connectivity strength in the semantic tasks (vs. the non-semantic task) for each pair of networks. Error bars indicate 95% confidence intervals.

#### Summary

Our network-level activation results indicate that semantic processing engages a broad range of semantic and domain-general networks. Moreover, ageing affects the recruitment of these networks. We found that older people activated DMN more than young people during semantic tasks and showed less down-regulation of DMN activity when task demands increased. In contrast, young people favoured the domain-general control network (i.e., MDN) when making semantic decisions and showed more sensitivity to increased task demands in this network than their older counterparts (though only for non-semantic processing). Ageing had no effects on the operation of the semantic-specific control network (SCN). We found no consistent age differences in functional connectivity between networks. These results support the DECHA model’s proposal of an age-related shift in reliance from executive functions to prior knowledge. However, our results also indicate that DECHA’s predictions of declined executive recruitment and flexibility in DMN-executive connectivity in older age may not apply to all cognitive domains. In the present study, the neural systems supporting semantic control were maintained in older age, which mirrored the intact semantic cognition abilities of older participants.

## Discussion

Age-related right hemisphere over activation is frequently reported but its underlying causes are still under debate. Compensation theories claim that increased right hemisphere engagement is a response to increased processing demands, while the dedifferentiation account suggests it is a result of age-related loss of specialisation and does not contribute to task performance. Our pre-registered fMRI study investigated this phenomenon within the domain of semantic cognition. Using knowledge- and control-demanding semantic tasks, we compared the effects of task demands on the activation of two core semantic regions – IFG and ATL – in the junior and senior brains. Consistent with the compensation view, we found higher task demands elicited stronger activation in right as well as left IFGs, across age groups and semantic tasks. This difficulty effect was also reproduced in the left ATL in the knowledge task, but not in the right ATL. However, we did not find age-related changes in the laterality of activation in either IFG or ATL. In addition to examining the compensation and dedifferentiation accounts, we also explored a new theory of cognitive ageing – DECHA (Spreng & Turner, 2019; Turner & Spreng, 2015). Consistent with the general scheme of DECHA, which suggests an age-related shifting reliance from cognitive control to prior knowledge in performing cognitive tasks, we found more engagement of DMN and less activation of MDN in the older people. We also found that older adults were worse at up-regulating their MDN activation as a function of difficulty in the non-semantic task (but not the semantic tasks). In this Discussion, we first consider effects in our pre-registered regions of interest in IFG and ATL, before turning to the analyses of large-scale brain networks.

### Responses to semantic task demands in IFGs and ATLs

Our key aim was to understand the true status of right hemisphere prefrontal activation in the domain of semantic cognition and its change with age. We found that both IFGs exhibited increased activation as a function of semantic demands, across age groups and semantic tasks. The effect in right IFG is in agreement with earlier meta-analyses, which tested effects of semantic control demands and found that right IFG showed higher activation for more controlled semantic tasks in young people (Jackson, 2021; Noonan, Jefferies, Visser, & Lambon Ralph, 2013). Our results establish that this demand-activation relationship in IFGs is present across age groups and in both knowledge- and control-demanding tasks. The right IFG increased its activation as both tasks became more difficult, which was accompanied with a similar but stronger difficulty effect in left IFG. This suggests that left IFG acts as the core frontal region supporting verbal semantics and that right IFG plays a more supporting role as task demands increase. Together, it appears that the left and right IFGs work in concert to regulate semantic processing in a task-demand-sensitive fashion, irrespective of age. These data support a compensation account of right IFG function, in which right IFG is recruited to help left IFG in task processing under more demanding situations. In contrast, it does not support the view that age-related over-activation of right IFG is due to neural dedifferentiation, as this effect would be independent of task demands.

Although our difficulty-based analyses supported the compensation view of right IFG recruitment, there was no age differences in activation of this region. Based on the compensation account, we predicted a higher activation in the right IFG in young participants in the semantic knowledge task, which was more demanding for them than the older population, and a reverse effect in the semantic control task. However, our results revealed a similar activation pattern for both age groups across tasks. This may have occurred because the average task difficulty in the current study was not high enough. According to compensation theories, age differences in right IFG should be most pronounced when task demands are high enough to push the left hemisphere beyond its processing limits in one age group (Reuter-Lorenz & Cappell, 2008). In our study, older people did perform significantly more accurately than young people on the knowledge task, but the difference between groups was relatively small (∼2%) and overall accuracy on both tasks was high (∼93%). It is therefore possible that the semantic tasks were not demanding enough to produce reliable age differences in core semantic regions (though we did find age effects at the network level, as discussed below). One important direction for future studies is to explore the IFG’s demand-activation profile with more challenging tasks and trials.

In contrast to right IFG, our study provided no evidence that the right ATL plays a compensatory role during semantic processing. Models of semantic cognition propose that the ATLs act as a hub for semantic knowledge representation (Jefferies, 2013; Lambon Ralph et al., 2017). In our study, the left ATL showed an age-independent difficulty effect in the knowledge task, consistent with this role. This difficulty effect was not replicated in the semantic control task, suggesting that left ATL is sensitive to knowledge representations but not to control demands. However, activation in the right ATL exhibited no relationship with semantic demands whatsoever. This seems contrary to an earlier computational modelling study proposing an integrated bilateral system of the left and right ATLs (Schapiro, McClelland, Welbourne, Rogers, & Lambon Ralph, 2013). Based on this model, failure in knowledge representation in one ATL can be compensated by an increased engagement of the other. This collaborating bilateral system is compatible with empirical findings from neuropsychological studies indicating that unilateral ATL damage only produces minor effects on semantic cognition (Lambon Ralph, Cipolotti, Manes, & Patterson, 2010; Lambon Ralph, Ehsan, Baker, & Rogers, 2012; Rice, Caswell, Moore, Hoffman, & Lambon Ralph, 2018).

Why then did we not observe right ATL engagement in our tasks? One possibility is that right ATL is only recruited when there is an impairment to the left ATL. If similar knowledge is encoded in both ATLs, there is no advantage to recruiting the right ATL when the left ATL is functioning normally – as all available knowledge can be retrieved through unilateral activation. In contrast, engagement of right ATL would be helpful when the information stored in the left ATL is degraded (e.g., by a disease process). This proposal is supported by a study that combined transcranial magnetic stimulation (TMS) with fMRI (Jung & Lambon Ralph, 2016). The researchers found that increased activity in right ATL predicted semantic performance after the left ATL had been inhibited by stimulation, but there was no correlation following stimulation of a non-semantic site. Another relevant factor is that knowledge for written word information is more strongly represented or easier to access in the left ATL. A left-lateralised bias in ATL for single written words has been found in numerous fMRI studies (Hoffman & Lambon Ralph, 2018; Rice, Hoffman, et al., 2015; Rice, Lambon Ralph, et al., 2015) and is consistent with the pattern of impairments in patients with left vs. right ATL resection (Rice, Caswell, Moore, Hoffman, et al., 2018). These findings indicate that input modality could also influence compensation effects in the right ATL. While the present verbal semantic tasks are heavily reliant on left ATL, a different picture could emerge for non-verbal stimuli.

The divergent activation profiles of the IFG and ATL also have implications for understanding their contributions to semantic processing. The Controlled Semantic Cognition framework proposes that semantic cognition is supported by the interaction of the semantic control network, with IFG as a critical area, and semantic knowledge network with ATL as its hub (Jefferies, 2013; Lambon Ralph et al., 2017). Consistent with this framework, we found that the IFG was more involved in the task emphasizing controlled semantic processing, while the ATL was more activated in the task that placed high demands on stored knowledge. However, these differences were fairly subtle and the two regions were robustly engaged in both semantic tasks. At the whole-brain level (Figure 5), the two semantic tasks also elicited largely similar areas of activation. These results suggest that regions specialised for semantic control and knowledge representation are highly interactive and cooperative in healthy participants, contributing to a variety of semantic tasks. This co-existence of specialisation and collaboration in the semantic system has also been demonstrated in neuropsychological studies. Different patient groups can fail the same semantic tasks for different underlying reasons (i.e., control vs. knowledge impairments), suggesting that semantic tasks rely on close interaction between these different systems (Jefferies & Lambon Ralph, 2006; Rogers, Patterson, Jefferies, & Lambon Ralph, 2015; Warrington & Cipolotti, 1996). The results of the present study also suggest a high level of overlap in the regions recruited for different types of semantic task.

### Network-level effects

Beyond IFG and ATL, the present study also investigated neurocognitive ageing effects at a network level. We included three networks of interest, among which the DMN has been linked with the representation of internal knowledge and experiences (Binder & Desai, 2011; Binder et al., 2009; Binder et al., 1999), and the MDN and SCN are related to domain-general and semantic-specific executive control, respectively (Fedorenko et al., 2013; Jackson, 2021). The choice of these networks was motivated by the DECHA hypothesis, built on the phenomenon that older people show losses of cognitive control and gains in crystallized knowledge (Spreng & Turner, 2019; Turner & Spreng, 2015). From a functional activation perspective, DECHA implies more engagement of the knowledge network (i.e., the DMN) and less activation of the executive control regions (e.g., the MDN and SCN) in older adulthood. Consistent with this prediction, our network-level activation results showed that the DMN was more activated for the semantic tasks in older people than the young, but the MDN was less activated. Interestingly, although both the MDN and SCN are linked with cognitive control (and both exhibited a preference for the semantic control task over the knowledge task), age-related decreases were only seen in MDN. These results support DECHA’s claim that there is an age-related shifting reliance from control ability to stored knowledge in supporting goal-directed behaviour. However, this shift appears only apply to general executive control regions but not semantic-specialised control regions.

Another observation relevant to DECHA is that older adults are less able to modulate activation of cognitive control areas in response to task demand (Cappell et al., 2010; Turner & Spreng, 2015). Our difficulty-based analyses investigated this. We found that MDN was less modulated by difficulty in older adults in the non-semantic task, but not in the semantic tasks. Moreover, no age difference in difficulty effect was found in the SCN. Thus, SCN regions did not show the same age-related changes as MDN. This is important because DECHA’s proponents focus on lateral prefrontal regions as the key site for cognitive control (Spreng & Turner, 2019), but this region contains both domain-general executive areas in MDN and semantic-specialised SCN regions (Jackson, 2021). Our results suggest that age-related changes apply mainly to the domain-general parts of this area and not to the more specialised SCN.

Besides the control regions, DECHA also predicts that older people are less able to supress DMN activation in the context of increasing task demands, which reflects their diminished ability to inhibit the influence of their prior knowledge. Our results showed that older people indeed tended to suppress DMN activation less than the young but this effect was mainly driven by the non-semantic task. In semantic tasks, the DMN may be making a positive contribution to performance, especially in older people. This is consistent with the task-based activation results, in which DMN was more active in the semantic tasks than non-semantic task in both groups.

From a functional connectivity perspective, DECHA proposes that older adults are worse at task-dependent modulation of the functional connectivity between knowledge and executive-control networks (Spreng & Turner, 2019; Turner & Spreng, 2015). On this view, the DMN and control networks exhibit positive coupling in all age groups when prior knowledge is congruent with task goals. However, when prior knowledge is not helpful or needs to be inhibited, only young people can decouple the DMN and control networks successfully. In our study, we found that DMN positively coupled with MDN and SCN during semantic tasks independent of age, confirming the importance of the interaction between knowledge and control networks for semantic processing (Martin, Saur, & Hartwigsen, 2022). Nevertheless, the predicted less-flexible DMN-executive connectivity in older age was not found: the difference in connectivity between semantic and non-semantic tasks was similar for both age groups. We do not have an obvious explanation for this. One possibility is that our non-semantic letter-matching task was so novel and perceptual in nature that neither age group attempted to engage their prior knowledge to complete it. It is worth noting that very few studies have investigated DMN-executive coupling in a similar context as our research (for exceptions, see Adnan et al., 2019; Amer, Giovanello, Nichol, Hasher, & Grady, 2019) and we believe future investigation on a wider range of tasks would benefit the field greatly.

In conclusion, by focusing on the domain of semantic cognition, our findings have provided a new perspective on the shifting architecture of cognition and brain function in later life. Our results revealed an age-independent compensatory mechanism in right IFG for semantic processing across age groups. At a large-scale network level, we found that older people engaged DMN more and MDN less than young people during semantic tasks, but showed no activation changes in the semantic-specific SCN. They also showed less demand-induced modulation in MDN, but only during non-semantic processing. These results suggest that semantic cognition can be relatively preserved, at the level of neural function as well as behaviour. This age-invariance in specialised semantic regions may help to explain the maintained performance of older people in semantic cognition.

## Acknowledgements

This work was supported by a BBSRC grant to P.H. (BB/T004444/1). Imaging was carried out at the Edinburgh Imaging Facility (www.ed.ac.uk/edinburgh-imaging), University of Edinburgh, which is part of the SINAPSE collaboration (www.sinapse.ac.uk). We are grateful to the University of Minnesota Center for Magnetic Resonance Research for sharing their neuroimaging sequences. We thank Yueyang Zhang for his help with data collection. We are also grateful to all research participants in this study. For the purpose of open access, the authors have applied a Creative Commons Attribution (CC BY) licence to any Author Accepted Manuscript version arising from this submission.

## Conflict of Interest

None declared.

## CRediT authorship contribution statement

Wei Wu: Conceptualization, Investigation, Formal analysis, Validation, Writing - original draft, Writing - review & editing, Visualization. Paul Hoffman: Conceptualization, Writing - review & editing, Project administration, Supervision, Funding acquisition.

## Notes

https://doi.org/10.7488/ds/3845

https://doi.org/10.7488/ds/3846

### Competing Interest Statement

The authors have declared no competing interest.

