## Supplementary Materials for "Age-related changes in the neural processing of semantics, within and beyond the core semantic network"

**Supplementary Information**

**Overview**

| Supplementary Figure S1 | Results of whole-brain univariate activation analysis (one-sample t test). |
| --- | --- |
| Supplementary Table S1 | Mixed effects models predicting accuracies and RTs in young and older people. |
| Supplementary Table S2 | Mixed effects models examining the shifting division of labour between IFG and ATL in different semantic tasks and age groups. |
| Supplementary Table S3 | Mixed effects models predicting ROI-level activation in the semantic tasks (vs. fixation) from age, task and hemisphere. |
| Supplementary Table S4 | Mixed effects models predicting ROI-level activation (vs. difficulty level 1) in the semantic tasks from age, difficulty and hemisphere. |
| Supplementary Table S5 | Correlations between behavioural performance score in each task and neural activation in each ROI for each age group. |
| Supplementary Table S6 | Mixed effects models predicting network-level activation (vs. the non-semantic task) from age and task. |
| Supplementary Table S7 | Mixed effects models predicting network-level activation (vs. difficulty level 1) from age, task and difficulty. |
| Supplementary Table S8 | Mixed effects models predicting network-level PPI functional connectivity changes (vs. the non-semantic task) from age and task. |

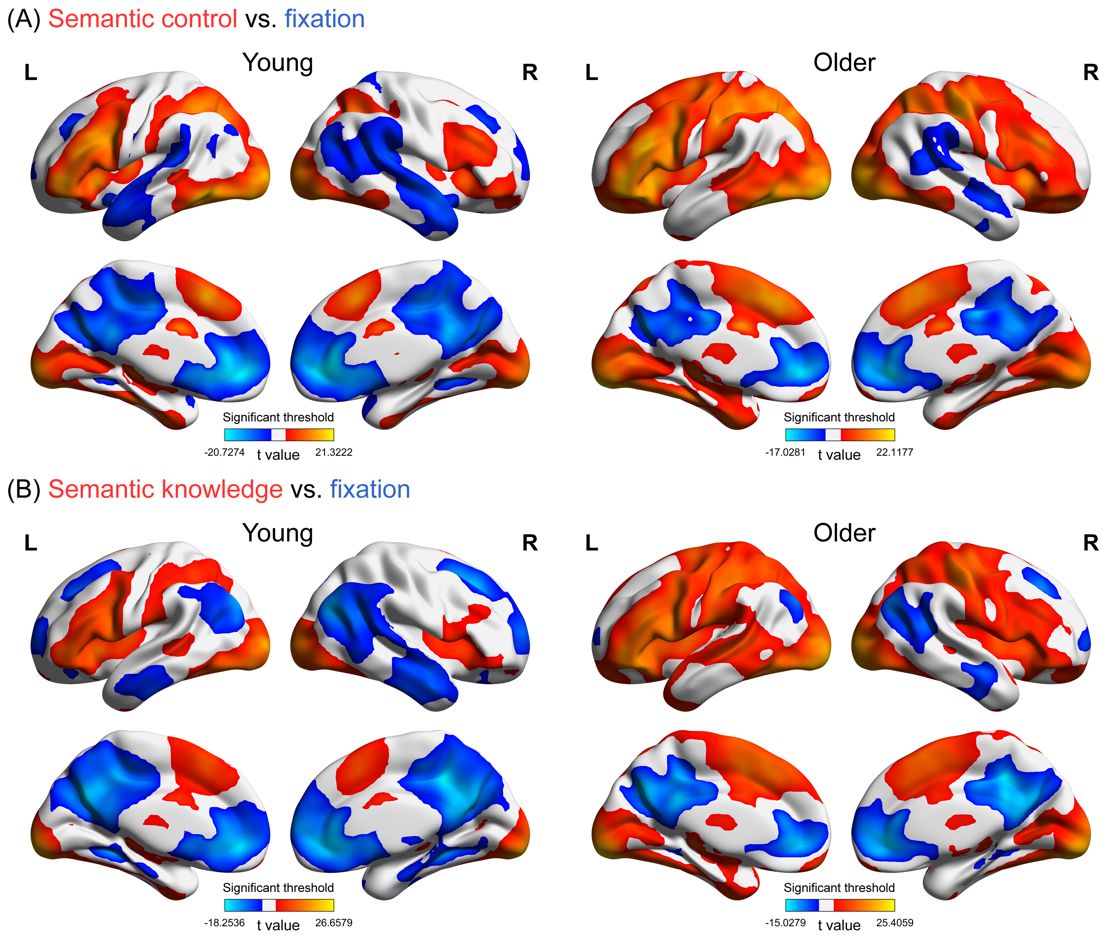

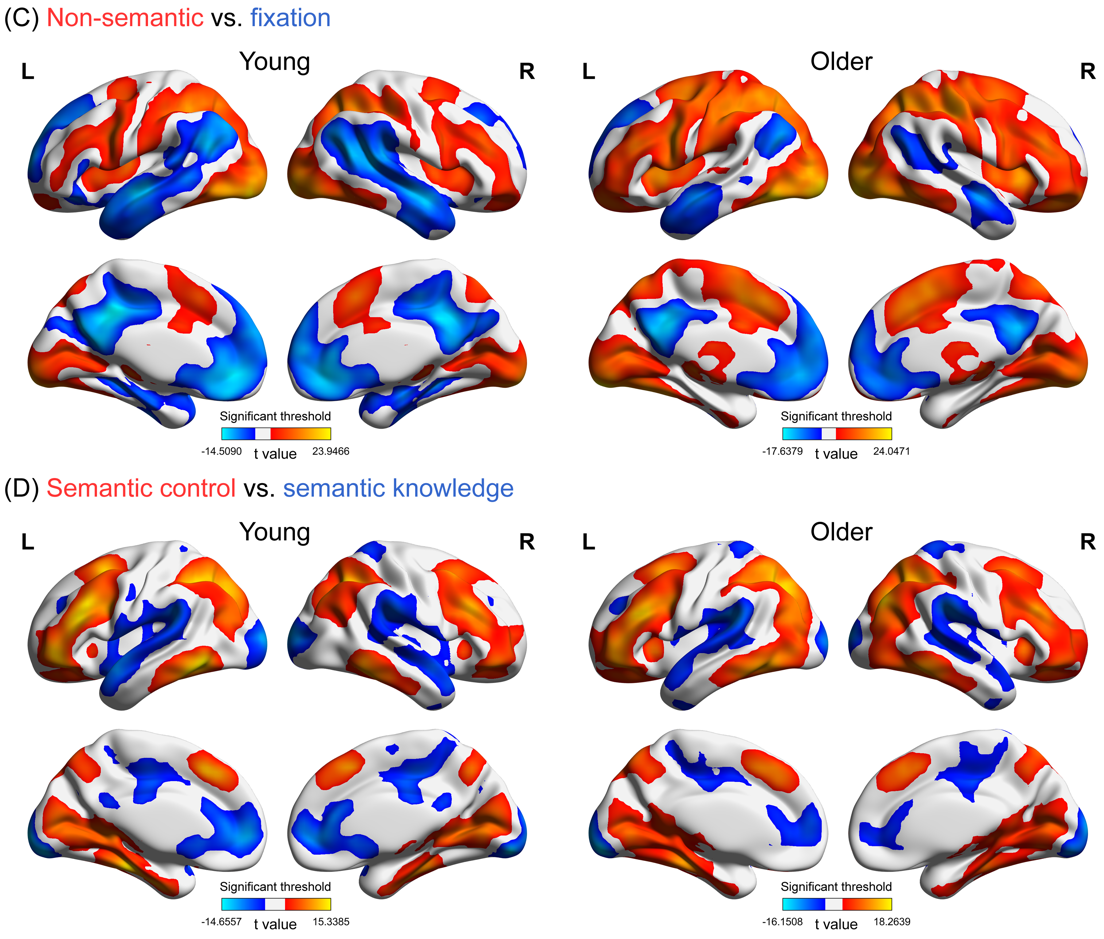

**Figure S1. Results of whole-brain univariate activation analysis (one-sample t test). Results were corrected for multiple comparisons, voxelwise p < 0.005, FWE corrected cluster threshold p = 0.05.**

**Table S1. Mixed effects models predicting accuracies and RTs in young and older people.**

|  | Accuracy | | |  | RT | | |
| --- | --- | --- | --- | --- | --- | --- | --- |
| Effect | B/Chisq | *s.e.* | *p* |  | B/F value | *s.e.* | *p* |
| *Age, task and difficulty effects* |  |  |  |  |  |  |  |
| Age | 0.693 | -- | 0.405 |  | 47.502 | -- | <10^-9^ |
| Task | 22.483 | -- | <10^-4^ |  | 58.059 | -- | <10^-15^ |
| Difficulty | 191.139 | -- | <10^-15^ |  | 356.329 | -- | <10^-15^ |
| Age × Task | 7.877 | -- | <0.05 |  | 13.541 | -- | <10^-5^ |
| Age × Difficulty | 1.788 | -- | 0.181 |  | 5.097 | -- | <0.05 |
| Task × Difficulty | 7.412 | -- | <0.05 |  | 4.756 | -- | <0.01 |
| Age × Task × Difficulty | 3.131 | -- | 0.209 |  | 4.022 | -- | <0.05 |
| *Age and difficulty effects in the semantic control task* |  |  |  |  |  |  |  |
| Age | 0.038 | 0.103 | 0.708 |  | 0.042 | 0.007 | <10^-7^ |
| Difficulty | -1.030 | 0.096 | <10^-15^ |  | 0.045 | 0.004 | <10^-15^ |
| Age × Difficulty | -0.112 | 0.073 | 0.117 |  | -0.001 | 0.002 | 0.750 |
| *Age and difficulty effects in the semantic knowledge task* |  |  |  |  |  |  |  |
| Age | 0.344 | 0.134 | <0.01 |  | 0.038 | 0.009 | <10^-4^ |
| Difficulty | -1.344 | 0.132 | <10^-15^ |  | 0.057 | 0.004 | <10^-15^ |
| Age × Difficulty | -0.166 | 0.111 | 0.065 |  | -0.001 | 0.003 | 0.658 |
| *Age and difficulty effects in the non-semantic task* |  |  |  |  |  |  |  |
| Age | -0.085 | 0.145 | 0.426 |  | 0.075 | 0.009 | <10^-11^ |
| Difficulty | -1.472 | 0.129 | <10^-15^ |  | 0.066 | 0.004 | <10^-15^ |
| Age × Difficulty | 0.022 | 0.104 | 0.719 |  | -0.009 | 0.002 | <0.001 |

Note: As the full accuracy model of the semantic control task failed to converge, we used a simplified model to omit random correlations from this model.

**Table S2. Mixed effects models examining the shifting division of labour between IFG and ATL in different semantic tasks and age groups.**

| Effect | B | *s.e.* | *p* |
| --- | --- | --- | --- |
| *Age, task and ROI effects* |  |  |  |
| Age | 0.017 | 0.022 | 0.443 |
| ROI | -0.307 | 0.010 | <10^-15^ |
| Task | 0.034 | 0.010 | <0.001 |
| Age × ROI | -0.018 | 0.010 | 0.066 |
| Age × Task | -0.014 | 0.010 | 0.161 |
| ROI × Task | -0.044 | 0.010 | <10^-4^ |
| Age × ROI × Task | 0.013 | 0.010 | 0.196 |

**Table S3. Mixed effects models predicting ROI-level activation in the semantic tasks (vs. fixation) from age, task and hemisphere.**

| Effect | B | *s.e.* | *p* |
| --- | --- | --- | --- |
| *Age, task and hemisphere effects in IFG* |  |  |  |
| Age | 0.035 | 0.032 | 0.269 |
| Hemisphere | 0.321 | 0.015 | <10^-15^ |
| Task | 0.078 | 0.015 | <10^-6^ |
| Age × Hemisphere | 0.016 | 0.015 | 0.290 |
| Age × Task | -0.027 | 0.015 | 0.087 |
| Hemisphere × Task | 0.032 | 0.015 | <0.05 |
| Age × Hemisphere × Task | -0.003 | 0.015 | 0.866 |
| *Age, task and hemisphere effects in ATL* |  |  |  |
| Age | -0.001 | 0.019 | 0.950 |
| Hemisphere | 0.076 | 0.007 | <10^-15^ |
| Task | -0.010 | 0.007 | 0.166 |
| Age × Hemisphere | 0.011 | 0.007 | 0.148 |
| Age × Task | -0.001 | 0.007 | 0.882 |
| Hemisphere × Task | -0.009 | 0.007 | 0.203 |
| Age × Hemisphere × Task | 0.003 | 0.007 | 0.673 |

**Table S4. Mixed effects models predicting ROI-level activation (vs. difficulty level 1) in the semantic tasks from age, difficulty and hemisphere.**

|  | IFG | | |  | ATL | | |
| --- | --- | --- | --- | --- | --- | --- | --- |
| Effect | B | *s.e.* | *p* |  | B | *s.e.* | *P* |
| *Control task* |  |  |  |  |  |  |  |
| Age | -0.096 | 0.034 | <0.01 |  | -0.015 | 0.022 | 0.496 |
| Hemisphere | 0.057 | 0.014 | <10^-4^ |  | 0.022 | 0.007 | <0.01 |
| Difficulty | 0.057 | 0.017 | <0.001 |  | -0.016 | 0.011 | 0.147 |
| Age × Hemisphere | -0.012 | 0.014 | 0.406 |  | -0.003 | 0.007 | 0.677 |
| Age × Difficulty | -0.0003 | 0.017 | 0.986 |  | -0.002 | 0.011 | 0.862 |
| Hemisphere × Difficulty | 0.018 | 0.011 | 0.097 |  | 0.005 | 0.007 | 0.479 |
| Age × Hemisphere × Difficulty | 0.007 | 0.011 | 0.536 |  | 0.006 | 0.007 | 0.376 |
| *Knowledge task* |  |  |  |  |  |  |  |
| Age | -0.130 | 0.039 | <0.01 |  | -0.039 | 0.022 | 0.082 |
| Hemisphere | 0.155 | 0.020 | <10^-10^ |  | 0.026 | 0.009 | <0.01 |
| Difficulty | 0.139 | 0.017 | <10^-11^ |  | 0.025 | 0.010 | <0.05 |
| Age × Hemisphere | -0.030 | 0.020 | 0.142 |  | -0.022 | 0.009 | <0.05 |
| Age × Difficulty | -0.003 | 0.017 | 0.867 |  | 0.006 | 0.010 | 0.560 |
| Hemisphere × Difficulty | 0.041 | 0.012 | <0.001 |  | 0.026 | 0.007 | <10^-4^ |
| Age × Hemisphere × Difficulty | 0.001 | 0.012 | 0.951 |  | -0.004 | 0.007 | 0.557 |
| *Control task, left hemisphere* |  |  |  |  |  |  |  |
| Age | -0.108 | 0.040 | <0.01 |  | -0.018 | 0.024 | 0.453 |
| Difficulty | 0.075 | 0.019 | <0.001 |  | -0.011 | 0.013 | 0.393 |
| Age × Difficulty | 0.006 | 0.019 | 0.741 |  | 0.004 | 0.013 | 0.729 |
| *Control task, right hemisphere* |  |  |  |  |  |  |  |
| Age | -0.084 | 0.033 | <0.05 |  | -0.012 | 0.022 | 0.591 |
| Difficulty | 0.039 | 0.017 | <0.05 |  | -0.021 | 0.011 | 0.054 |
| Age × Difficulty | -0.007 | 0.017 | 0.683 |  | -0.008 | 0.011 | 0.445 |
| *Knowledge task, left hemisphere* |  |  |  |  |  |  |  |
| Age | -0.160 | 0.047 | <0.01 |  | -0.061 | 0.025 | <0.05 |
| Difficulty | 0.180 | 0.022 | <10^-11^ |  | 0.052 | 0.012 | <10^-4^ |
| Age × Difficulty | -0.002 | 0.022 | 0.926 |  | 0.002 | 0.012 | 0.856 |
| *Knowledge task, right hemisphere* |  |  |  |  |  |  |  |
| Age | -0.099 | 0.040 | <0.05 |  | -0.016 | 0.022 | 0.469 |
| Difficulty | 0.098 | 0.015 | <10^-8^ |  | -0.001 | 0.010 | 0.919 |
| Age × Difficulty | -0.004 | 0.015 | 0.809 |  | 0.010 | 0.010 | 0.326 |

Note: As the right IFG model of the semantic knowledge task failed to converge, we used a simplified model to omit random correlations from this model.

**Table S5. Correlations between behavioural performance score in each task and neural activation in each ROI for each age group.**

|  | *r* value (*p* value) | | | |
| --- | --- | --- | --- | --- |
|  | Left IFG | Right IFG | Left ATL | Right ATL |
| *Older* |  |  |  |  |
| Control task | 0.30 (0.13) | 0.25 (0.16) | 0.40 (<0.05) | 0.27 (0.15) |
| Knowledge task | 0.17 (0.34) | 0.11 (0.55) | 0.27 (0.15) | 0.33 (0.09) |
| Non-semantic task | 0.09 (0.57) | 0.18 (0.32) | 0.48 (<0.05) | 0.34 (0.09) |
| *Young* |  |  |  |  |
| Control task | -0.27 (0.33) | 0.08 (0.89) | 0.11 (0.80) | 0.28 (0.33) |
| Knowledge task | -0.27 (0.33) | 0.06 (0.89) | -0.18 (0.60) | 0.13 (0.80) |
| Non-semantic task | 0.02 (0.97) | 0.05 (0.89) | -0.24 (0.35) | 0.01 (0.97) |

Note: The p values are FDR-corrected for multiple comparisons.

**Table S6. Mixed effects models predicting network-level activation (vs. the non-semantic task) from age and task.**

| Effect | B | *s.e.* | *p* |
| --- | --- | --- | --- |
| *MDN* |  |  |  |
| Age | -0.187 | 0.044 | <10^-4^ |
| Task | 0.249 | 0.011 | <10^-15^ |
| Age × Task | 0.018 | 0.011 | 0.119 |
| *SCN* |  |  |  |
| Age | -0.060 | 0.039 | 0.133 |
| Task | 0.259 | 0.013 | <10^-15^ |
| Age × Task | 0.003 | 0.013 | 0.840 |
| *DMN* |  |  |  |
| Age | 0.312 | 0.039 | <10^-11^ |
| Task | -0.013 | 0.013 | 0.300 |
| Age × Task | -0.003 | 0.013 | 0.822 |

**Table S7. Mixed effects models predicting network-level activation (vs. difficulty level 1) from age, task and difficulty.**

| Effect | F value | *p* |
| --- | --- | --- |
| *MDN* |  |  |
| Age | 9.664 | <0.01 |
| Task | 64.304 | <10^-15^ |
| Difficulty | 561.619 | <10^-15^ |
| Age × Task | 8.108 | <0.001 |
| Age × Difficulty | 15.915 | <0.001 |
| Task × Difficulty | 99.350 | <10^-15^ |
| Age × Task × Difficulty | 10.163 | <10^-4^ |
| *SCN* |  |  |
| Age | 8.617 | <0.01 |
| Task | 22.820 | <10^-7^ |
| Difficulty | 586.215 | <10^-15^ |
| Age × Task | 0.942 | 0.394 |
| Age × Difficulty | 0.233 | 0.630 |
| Task × Difficulty | 1.270 | 0.286 |
| Age × Task × Difficulty | 0.738 | 0.481 |
| *DMN* |  |  |
| Age | 3.961 | <0.05 |
| Task | 45.156 | <10^-13^ |
| Difficulty | 67.978 | <10^-12^ |
| Age × Task | 3.976 | <0.05 |
| Age × Difficulty | 15.234 | <0.001 |
| Task × Difficulty | 34.490 | <10^-12^ |
| Age × Task × Difficulty | 0.859 | 0.426 |

**Table S8. Mixed effects models predicting network-level PPI functional connectivity changes (vs. the non-semantic task) from age and task.**

|  | MDN - DMN | | |  | SCN - DMN | | |  | MDN - SCN | | |
| --- | --- | --- | --- | --- | --- | --- | --- | --- | --- | --- | --- |
| Effect | B | *s.e.* | *p* |  | B | *s.e.* | *p* |  | B | *s.e.* | *p* |
| Age | -0.011 | 0.013 | 0.396 |  | -0.023 | 0.021 | 0.273 |  | 0.033 | 0.018 | 0.077 |
| Task | 0.046 | 0.010 | <10^-5^ |  | 0.047 | 0.011 | <10^-4^ |  | 0.046 | 0.008 | <10^-7^ |
| Age × Task | -0.004 | 0.010 | 0.661 |  | 0.013 | 0.011 | 0.261 |  | -0.019 | 0.008 | <0.05 |
